# Uremic toxin indoxyl sulfate induces trained immunity *via* the AhR-dependent arachidonic acid pathway in end-stage renal disease (ESRD)

**DOI:** 10.1101/2022.08.08.503143

**Authors:** Hee Young Kim, Yeon Jun Kang, Dong Hyun Kim, Jiyeon Jang, Su Jeong Lee, Gwanghun Kim, Hee Byung Koh, Ye Eun Ko, Hyun Mu Shin, Hajeong Lee, Tae-Hyun Yoo, Won-Woo Lee

## Abstract

Trained immunity is the long-term functional reprogramming of innate immune cells, which results in altered responses toward a secondary challenge. Despite indoxyl sulfate (IS) being a potent stimulus associated with chronic kidney disease (CKD)-related inflammation, its impact on trained immunity has not been explored. Here, we demonstrate that IS induces trained immunity in monocytes *via* epigenetic and metabolic reprogramming, resulting in augmented cytokine production. Mechanistically, the aryl hydrocarbon receptor (AhR) contributes to IS-trained immunity by enhancing the expression of arachidonic acid (AA) metabolism-related genes such as Arachidonate 5-Lipoxygenase (ALOX5) and ALOX5 Activating Protein (ALOX5AP). Inhibition of AhR during IS training suppresses the induction of IS-trained immunity. Monocytes from end-stage renal disease (ESRD) patients have increased ALOX5 expression and after 6-day training, they exhibit enhanced TNF-α and IL-6 production to LPS. Furthermore, healthy control-derived monocytes trained with uremic sera from ESRD patients exhibit increased production of TNF-α and IL-6. Consistently, IS-trained mice and their splenic myeloid cells had increased production of TNF-α after *in vivo* and *ex vivo* LPS stimulation compared to that of control mice. These results provide insight into the role of IS in the induction of trained immunity, which is critical during inflammatory immune responses in CKD patients.

## Introduction

Over the last decade, a large body of evidence has demonstrated that innate cells can build up immunological memory resulting in enhanced responsiveness to subsequent stimulation, a phenomenon termed trained immunity (1). Compared with classical epitope-specific adaptive immunological memory based on an antigen-receptor, trained immunity of monocytes and macrophages is the long-term functional reprogramming elicited by an initial primary insult, mainly pathogen-associated molecular patterns (PAMPs), which leads to an altered response towards a subsequent, unrelated secondary insult after the return to a homeostatic state (2, 3). It has been well demonstrated that exposure of monocytes or macrophages to *Candida albicans*, fungal cell wall component β-glucan, or BCG (Bacille Calmette-Guérin) vaccine enhances their subsequent responses to unrelated pathogens or pathogen components such as lipopolysaccharide (LPS) (4, 5). The induction of trained immunity is associated with the interaction of epigenetic modifications and metabolic rewiring, which can last for prolonged periods of time (2, 3, 5–8). Mechanistically, certain metabolites derived from the upregulation of different metabolic pathways triggered by primary insult can influence enzymes involved in remodeling the epigenetic landscape of cells. This leads to specific changes in epigenetic histone markers, such as histone 3 lysine 4 trimethylation (H3K4me3) or histone 3 lysine 27 acetylation (H3K27ac), which regulate genes resulting in a more rapid and stronger response upon a subsequent, unrelated secondary insult (2–4, 7). In addition, it has been recently reported that long non-coding RNAs induce epigenetic reprogramming *via* the histone methyltransferase, MLL1. Subsequently, transcription factors such as Runx1 regulate the induction of proinflammatory cytokines following the secondary insult (9–11).

Many studies have provided evidence that trained immunity likely evolved as a beneficial process for non-specific protection from future secondary infections (3). However, it has also been suggested that augmented immune responses resulting from trained immunity is potentially relevant to deleterious outcomes in immune-mediated and chronic inflammatory diseases such as autoimmune diseases, allergy, and atherosclerosis (8, 10, 12–15). Thus, although most studies have focused on the ability of exogenous microbial insults to induce trained immunity, it is also conceivable that sterile inflammatory insults can evoke trained immunity. In support of this idea, oxLDL, lipoprotein a (Lpa), uric acid, hyperglycemia, and the Western diet have all been recently identified as endogenous sterile insults that induce trained immunity in human monocytes *via* epigenetic reprogramming (8, 10, 12, 13, 16). Thus, it is tempting to speculate that many endogenous insults that cause chronic inflammatory conditions may be involved in the induction of trained immunity in human monocytes and macrophages.

Chronic kidney disease (CKD) is recognized as a major non-communicable disease with increasing worldwide prevalence (17, 18). Loss of renal function in CKD patients causes the accumulation of over 100 uremic toxins, which are closely associated with cardiovascular risk and mortality due to their ability to generate oxidative stress and a proinflammatory cytokine milieu (19). Reflecting this, cardiovascular disease (CVD) is a leading cause of death among patients with end-stage renal disease (ESRD) (20). Indoxyl sulfate (IS) is a major uremic toxin derived from dietary tryptophan *via* fermentation of gut microbiota (21). Since it is poorly cleared by hemodialysis, IS is one of the uremic toxins present at higher than normal concentrations in the serum of CKD patients (22, 23) and is associated with the progression of CKD and the development of CKD-related complications such as CVD (24). We and others have shown that IS promotes the production of proinflammatory cytokines such as TNF-α and IL-1β by monocytes and macrophages through aryl hydrocarbon receptor (AhR) signaling and organic anion transporting polypeptides 2B1 (OATP2B1)-Dll4-Notch Signaling (21, 25, 26), suggesting a role of IS as an endogenous inflammatory insult in monocytes and macrophages. Moreover, pretreatment with IS greatly increases TNF-α production by human macrophages in response to a low dose of LPS (25). Despite the function of IS as an endogenous inflammatory insult in monocytes and macrophages, little is known with regard to whether IS induces trained immunity. Thus, we investigated whether exposure to IS triggers trained immunity in an *in vitro* human monocyte model and an *in vivo* mouse model, as well as the mechanisms involved in IS-induced trained immunity. Our data show that IS triggers trained immunity in human monocytes/macrophages *via* AhR-dependent alteration of the arachidonic acid (AA) pathway, epigenetic modifications, and metabolic rewiring. Thus, this suggests IS plays a critical role in the initiation of inflammatory immune responses in patients with CKD.

## Results

### Indoxyl sulfate (IS) induces trained immunity in human monocytes

To explore whether exposure to IS is involved in the induction of trained immunity in human monocytes, an *in vitro* model of trained immunity was applied as previously reported by the Netea group (27). Freshly isolated human CD14^+^ monocytes were preincubated for 24 hrs with or without IS and, after a subsequent 5-day culture in human serum, restimulated with lipopolysaccharide (LPS) or Pam3cys for final 24 hrs (Fig. 1A). Preincubation of monocytes with IS led to enhanced production of TNF-α, a major monocyte/macrophage-derived inflammatory cytokine, upon LPS stimulation. Since 10 ng/ml of LPS significantly increased both TNF-α and IL-6 secretion in IS-trained macrophages (Fig. 1B), we used this concentration of LPS in subsequent experiments. A clinically relevant concentration of IS in severe CKD has reported the range from 0.5 to 1.0 mmol/L (19). The preincubation effect of IS on cytokine production was observed at a concentration as low as 250 μM, which is the average IS concentration in patients with ESRD in our cohort (Fig. 1C) (25). Unlike IS, preincubation with other protein-bound uremic toxins (PBUTs), such as *p*-cresyl sulfate (PCS), Hippuric acid (HA), Indole 3-acetic acid (IAA) and kynurenic acid (KA), did not cause increased secretion of TNF-α or IL-6 in response to LPS stimulation (Fig. S1A). In addition, there was no obvious effect on cell viability following pre-incubation of macrophages with 1,000 μM of IS after a subsequent 5-day culture in human serum or after LPS stimulation (Fig. S1B). We also found that the enhanced cytokine production of IS-trained macrophages was not attributable to potassium derived from IS potassium salt (Fig. S1C). Moreover, the increased TNF-α and IL-6 production in IS-trained macrophages was not limited to LPS stimulation, as similar phenomena were observed following stimulation with Pam3cys, a TLR1/2 agonist (Fig. 1D). β-glucan-pretreated macrophages exhibit a prototypic feature of trained immunity, characterized by enhanced production of inflammatory cytokines upon restimulation with heterologous stimuli, LPS or Pam3cys (6, 27). As seen in Figure S1D, the level of TNF-α secreted by IS-trained macrophages was comparable with that secreted by β-glucan-trained macrophages, although β-glucan had a more potent effect on IL-6 production than did IS, suggesting that IS plays a role in the induction of trained immunity of human monocytes. Furthermore, alongside elevated TNF-α and IL-6 expressions, there was a notable increase in the mRNA expression of *IL-1β* and *MCP-1* (*CCL2*) observed in IS-trained macrophages, concomitant with a significant reduction in *IL-10*, a cytokine known for its anti-inflammatory properties, within the same cellular context (Fig. 1E). Correspondingly, alterations in their protein levels mirrored the observed mRNA expressions (Fig. S1E). Circulating monocytes have been identified as a major immune cell subset that responds to IS in the serum of ESRD patients (21, 25). To examine whether uremic serum induces trained immunity of monocytes/macrophages, pooled sera from ESRD patients (184 ± 44 μM of average IS level) or from HCs were used to treat monocytes isolated from HCs for 24 hr at 30% (v/v), followed by training for 5 days (Fig. 1F). Training with pooled uremic serum of ESRD patients increased the production of TNF-α and IL-6 upon re-stimulation with LPS compared to monocytes treated with the pooled sera of HCs (Fig. 1G-I). In addition, expression of *IL-1β* and *MCP-1* mRNA was also augmented by training with the pooled uremic sera of ESRD patients (Fig. 1I). These results suggest that IS induces trained immunity in human monocytes, characterized by the increased expression of proinflammatory cytokines TNF-α and IL-6 and reduced expression of anti-inflammatory IL-10 in response to secondary TLR stimulation.

**Figure 1.**
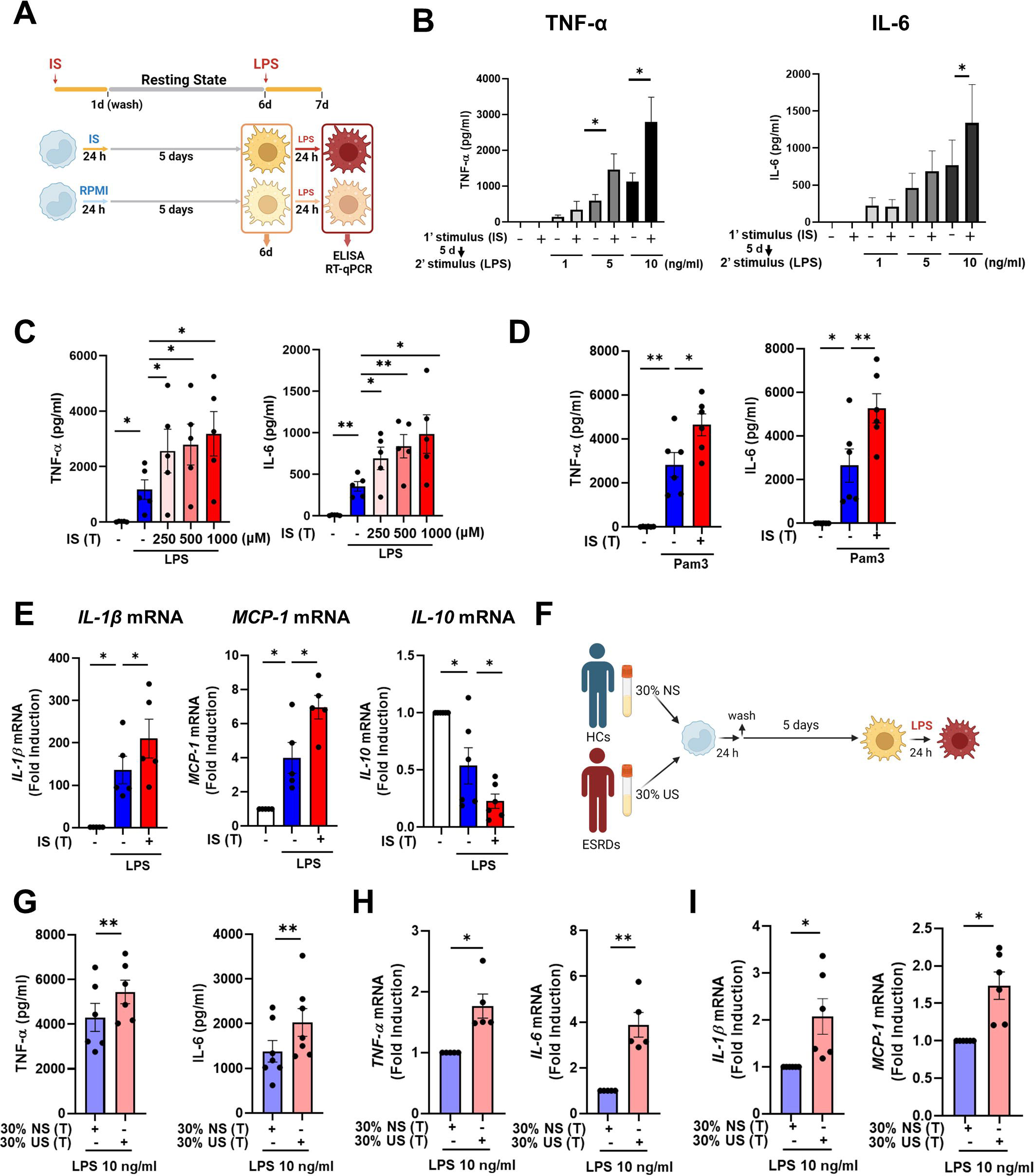
IS induces trained immunity in human monocytes. **A.** Schematic of *in vitro* experimental model for innate trained immunity. **B-C.** Human monocytes were treated with the indicated concentration of IS for 24 hr, followed by a subsequent 5-day culture in human serum. On day 6, the cells were restimulated with the indicated concentrations of LPS for 24 hr. TNF-α and IL-6 proteins levels were quantified by ELISA. **D.** After training with 1,000 μM IS, monocytes were restimulated with 10 μg/ml Pam3cys. TNF-α and IL-6 protein levels were quantified by ELISA. **E.** After training with 1,000 μM IS, monocytes were restimulated with 10 ng/ml LPS for 24 hr. The mRNA expression of *IL-1β*, *IL-10*, and *MCP-1* was analyzed by RT-qPCR. **F.** *In vitro* experimental scheme of uremic serum-induced trained immunity. **G-I**. The pooled normal serum (NS) from healthy controls (HCs) or uremic serum (US) from patients with ESRD were used for treatment of monocytes isolated from HCs for 24 hr at 30% (v/v) followed by resting for 5 days. After stimulation with LPS for 24 hr, TNF-α and IL-6 production were analyzed using ELISA (G) and RT-qPCR (H). After stimulation with LPS (10 ng/ml) for 24 hr, mRNA expression of *IL-1β* and *MCP-1* were determined by RT-qPCR (I). Bar graphs show the mean ± SEM. * = *p* < 0.05, and ** = *p* < 0.01 by two-tailed paired *t*-test.

### IS-induced trained immunity is regulated by metabolic rewiring

Metabolic rewiring is one of the most crucial processes regulating the trained immunity of monocytes and macrophages (33). Assessment of the metabolic profile of IS-trained macrophages on day 6 (prior to re-stimulation with LPS) showed that training with IS led to an enhanced extracellular acidification rate (ECAR) as a measure of lactate production, indicating increased glycolysis and glycolysis capacity (Fig. 2A and B). Moreover, basal and maximal respiration and ATP production gauged by the oxygen consumption rate (OCR) were also increased compared to that of non-trained cells (Fig. 2C and D). Enhanced glycolysis and glycolytic capacity in IS-trained cells remained higher even after re-stimulation with LPS (Fig. S2A-B), implying that the IS-training effect on metabolic rewiring is sustained regardless of the secondary stimulation. To further examine whether the metabolic rewiring by IS-trained cells is linked to the regulation of trained immunity, 2-deoxy-d-glucose (2-DG), a general inhibitor of glycolysis, was added to monocytes before training with IS. 2-DG completely inhibited the augmented production of TNF-α and IL-6 in IS-trained macrophages in response to re-stimulation with LPS (Fig. 2E). These data demonstrate that IS-trained immunity is linked to metabolic rewiring characterized by both enhanced glycolysis and augmented oxidative respiration.

**Figure 2.**
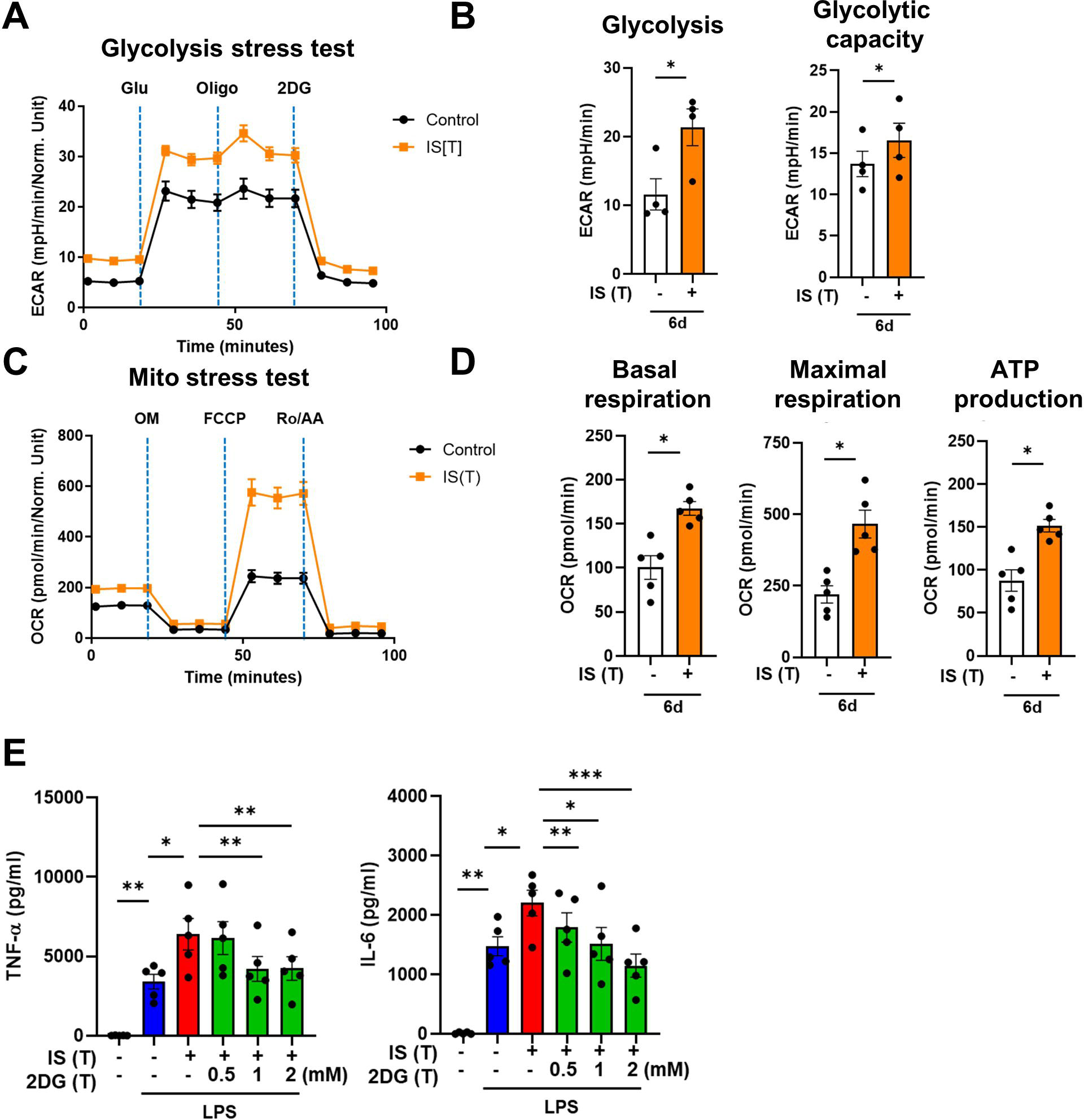
IS-induced trained immunity is linked to metabolic rewiring. Glycolysis and mitochondrial stress tests were conducted on IS (1,000 μM)-trained macrophages (n = 3∼4) using the Seahorse XF-analyzer. **A.** ECAR (extracellular acidification rate) levels were measured after sequential treatment with glucose, oligomycin, and 2-DG. **B.** Cellular glycolysis and glycolytic capacity were analyzed. **C.** OCR (Oxygen consumption rate) levels were measured after sequential treatment with oligomycin, FCCP, and Rotenone/antimycin A (Ro/AA). **D.** Basal respiration, maximal respiration, and ATP production were analyzed. **E.** Monocytes were pretreated with 2DG, followed by IS-training for 6 days. Cells were restimulated with LPS for 24 hr and TNF-α and IL-6 in supernatants were quantified by ELISA (n = 5). Bar graphs show the mean ± SEM. *= *p* < 0.05, **= *p* < 0.01, and *** = *p*< 0.001 by two-tailed paired *t*-test.

### Epigenetic modifications control IS-induced trained immunity

The induction of trained immunity relies on two key, closely intertwined mechanisms, epigenetic modification and metabolic rewiring of innate immune cells (2, 28, 29). We next sought to determine whether increased expression of TNF-α and IL-6 is a result of epigenetic changes. To this end, chromatin modification of histone 3 trimethylation of lysine 4 (H3K4me3) at the promoter sites of *TNFA* and *IL6* was analyzed. Chromatin immunoprecipitation (ChIP)-qPCR data illustrate that IS-trained macrophages exhibit enhanced H3K4me3 of *TNFA* and *IL6* promoters by day 6 after treatment with 1,000 μM of IS (Fig. 3A and B). This reflects what was previously demonstrated in trained innate immune cells (4, 22, 30). Moreover, IS-mediated enrichment of H3K4me3 was maintained even after secondary stimulation with LPS compared with non-trained cells (Fig. S3A-B). When IS-trained macrophages were pretreated with 5’- methylthioadenosine (MTA), a non-selective methyltransferase inhibitor, their production of TNF-α and IL-6 upon LPS stimulation was reversed to baseline (Fig. 3C), implying that IS-induced trained immunity is associated with epigenetic modification. To explore the potential regulation of IS-induced epigenetic modification by metabolic rewiring, we examined the enrichment of H3K4me3 at the promoters of *TNFA* and *IL6* subsequent to treatment with 2DG (Fig. 3D). Our findings suggest that metabolic rewiring influences epigenetic modification, implicating the participation of metabolites. Additionally, heightened enrichment of H3K4me3 at the promoter regions of *HK2* and *PFKP*, pivotal genes associated with glycolysis, was observed (Fig. S3C). To further elucidate epigenetic modifications in IS-induced trained immunity, we performed a whole-genome assessment of the histone marker H3K4me3 by ChIP-sequencing (ChIP-Seq) in IS-trained cells on day 6. Among 7,136 peaks, 59 differentially upregulated peaks and 316 downregulated peaks were detected in IS-trained cells (Fig. 3E and Table S1). To identify the biological processes affected in IS-mediated trained immunity, 59 upregulated peaks in IS-trained macrophages were analyzed through Gene Ontology (GO) analysis with Go biological process and the Reactome Gene Set. Activation of the innate immune response and positive regulation of the defense response were identified as major processes *via* Go biological process analysis. Further, genes involved in regulation of ornithine decarboxylase (ODC) and metabolism of polyamine were recognized as major gene sets *via* Reactome Gene Set analysis (Fig. 3F). A genome browser snapshot showing H3K4me3 binding illustrates that H3K4me3 is elevated at the promoters of important target genes associated with activation of the innate immune response, such as *IFI16* (interferon-gamma inducible protein 16), *XRCC5* (X-ray repair cross-complementing 5), and *PQBP1* (polyglutamine binding protein 1) and genes linked to the regulation of ornithine decarboxylase (ODC), such as *PSMA1* (proteasome 20S subunit alpha 1), *PSMA3* (proteasome 20S subunit alpha 3), and *OAZ3* (Ornithine Decarboxylase Antizyme 3, a protein that negatively regulates ODC activity) (Fig. 3G) (31). Additionally, differences in H3K4me3 enrichment patterns between the IS-training group and the control group were observed in *TNFA* and *IL6* (Fig. S3D-E). Our results show that epigenetic modification of innate immune response-related genes contributes to the induction of IS-trained immunity in human monocytes.

**Figure 3.**
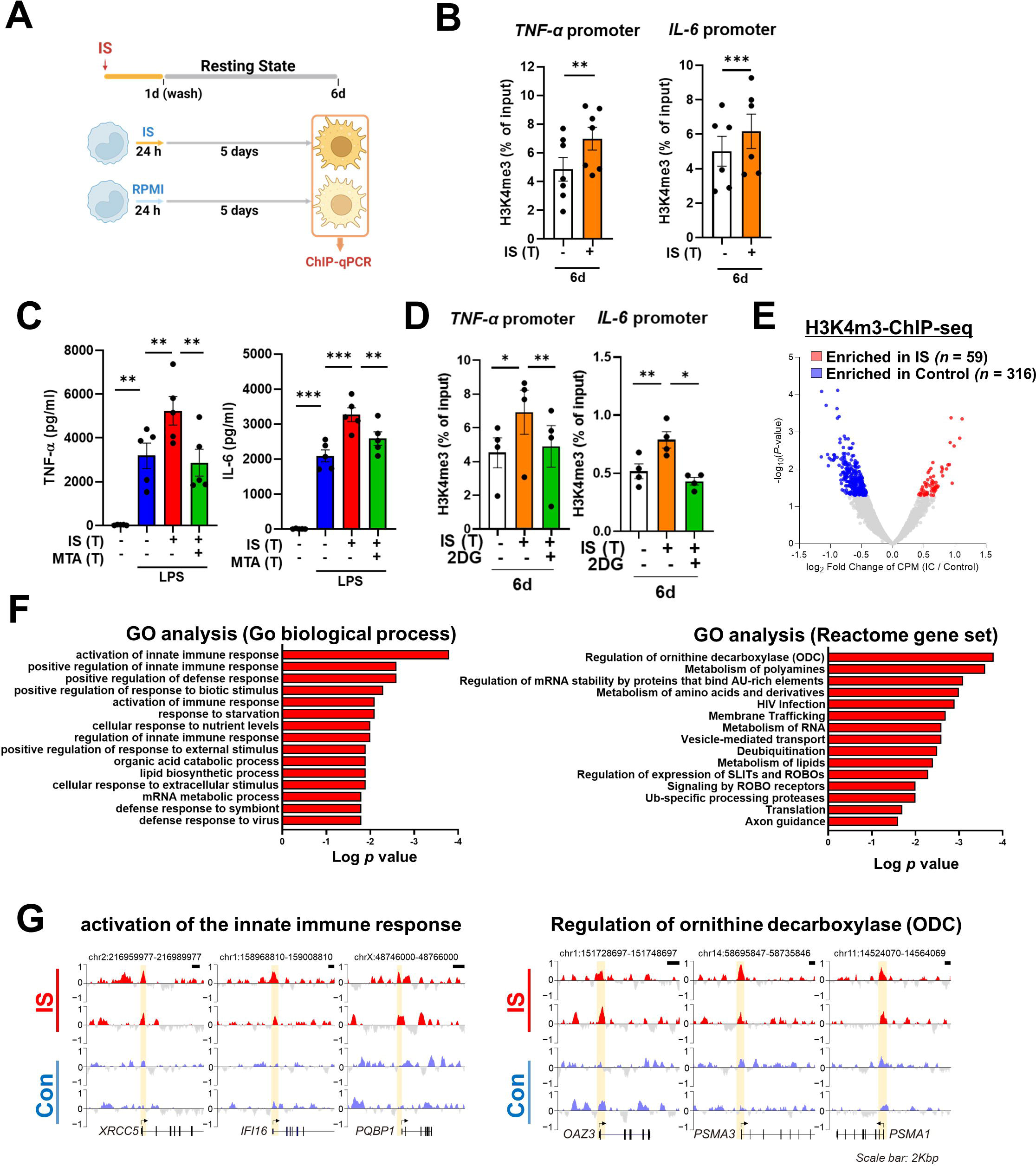
IS-induced trained immunity is accomplished through epigenetic modification. **A.** Experimental scheme of ChIP-qPCR for IS (1,000 μM)-trained macrophages. **B.** On day 6 after IS-training, cells were fixed with 1% formaldehyde, lysed, and sonicated. A ChIP assay was performed using anti-H3K4me3 antibody and enrichment of H3K4me3 in the promoter site of *TNFA* and *IL6* loci was quantified by qPCR. 1% input was used as a normalization control. **C.** Monocytes were pre-treated with 5’-methylthioadenosine (MTA, a non-selective methyltransferase inhibitor; 200 μM) and then were trained with IS for 6 days, followed by restimulation with LPS for 24 hrs. TNF-α and IL-6 proteins levels were quantified by ELISA. **D.** A ChIP assay was performed in IS-trained macrophages pre-treated with 2DG. 2% input was used as a normalization control. **E.** ChIP-Seq analysis was performed with anti-H3K4me3 antibody on chromatin isolated at day 6 from IS-trained and control macrophages. Enriched peaks in ChIP-Seq on H3K4me3 are shown as a volcano plot. (FC > 1.3, *p* < 0.05) **F.** Functional annotation of 59 upregulated Differentially regulated peaks (DRPs) on H3K4me3 in IS-trained macrophages were analyzed by Gene Ontology (Go) analysis with Go biological pathway and Reactome gene sets (FC > 1.3, *p* < 0.05). **G.** Screen shots of H3K4me3 modification in the promoter regions of *IFI16, XRCC5, PQBP1 PSMA1, PSMA3,* and *OAZ3*. *= *p*< 0.05, **= *p* < 0.01, and *** = *p* < 0.001 by two-tailed paired *t*-test.

### AhR, a potent endogenous receptor for IS, contributes to the induction of IS-trained immunity

Our previous study demonstrated that IS-induced TNF-α production in macrophages is regulated through a complex mechanism involving the interaction of NF-κB and SOCS2 with AhR (25). To explore the molecular mechanism underlying the regulation of IS-trained immunity, we investigated the role of AhR, a potent endogenous receptor for IS. Ligand-bound activated AhR is known to be immediately translocated into the nucleus and rapidly degraded (25, 32–34). Immunoblot analysis depicted in Figure 4A reveals persistent nuclear translocation of IS-mediated AhR even on day 6 (prior to re-stimulation with LPS), which was entirely inhibited by GNF351 treatment, an AhR antagonist, on day 6. Inhibition of AhR by GNF351 during IS training suppressed the increase in production of TNF-α and IL-6 following LPS restimulation on day 6 in IS-trained cells (Fig. 4B and C), implying that IS-mediated AhR activation may be involved in trained immunity. In addition to TNF-α and IL-6, enhancement of *IL-1β* and *MCP-1* mRNA expression in IS-trained cells was also completely inhibited, whereas decreased *IL-10* expression was completely reversed by GNF351 (Fig. 4D). To confirm the regulatory role of AhR in trained immunity, we tested whether 6-Formylindolo[3,2-b]carbazole (FICZ), a tryptophan-derived agonist of AhR, also induced trained immunity in human monocytes. FICZ-pretreated monocytes exhibited augmented expression of TNF-α and IL-6 in response to secondary stimulation with LPS compared to non-trained cells (Fig. S4A). Additionally, knockdown of AhR suppressed the expression of TNF-α and IL-6 in IS-trained cells (Fig. 4E), underscoring the significant role of ligand-bound activated AhR in the trained immunity of human monocytes. We next examined whether inhibition of AhR with GNF351 influences epigenetic modification and metabolic rewiring. Our ChIP-qPCR assay showed that enrichment of H3K4m3 on *TNFA* and *IL6* promoters in IS-trained macrophages was inhibited by GNF351 (Fig. 4F). Of note, assessment of the metabolic profile by measuring ECAR and OCR illustrates that GNF351 has no effect on metabolic rewiring, including enhanced glycolysis and mitochondrial respiration, in IS-trained cells on day 6 as depicted in Figure 3 (Fig. S4B-C). This finding was corroborated by the immunoblotting data, which showed GNF351 had no inhibitory effect on IS-mediated enhancement of S6K activity, which is critical for inducing the aerobic glycolysis in human monocytes/macrophages (Fig. S4D). Our findings suggest that IS-activated AhR is involved in regulating epigenetic modifications of IS-trained macrophages.

**Figure 4.**
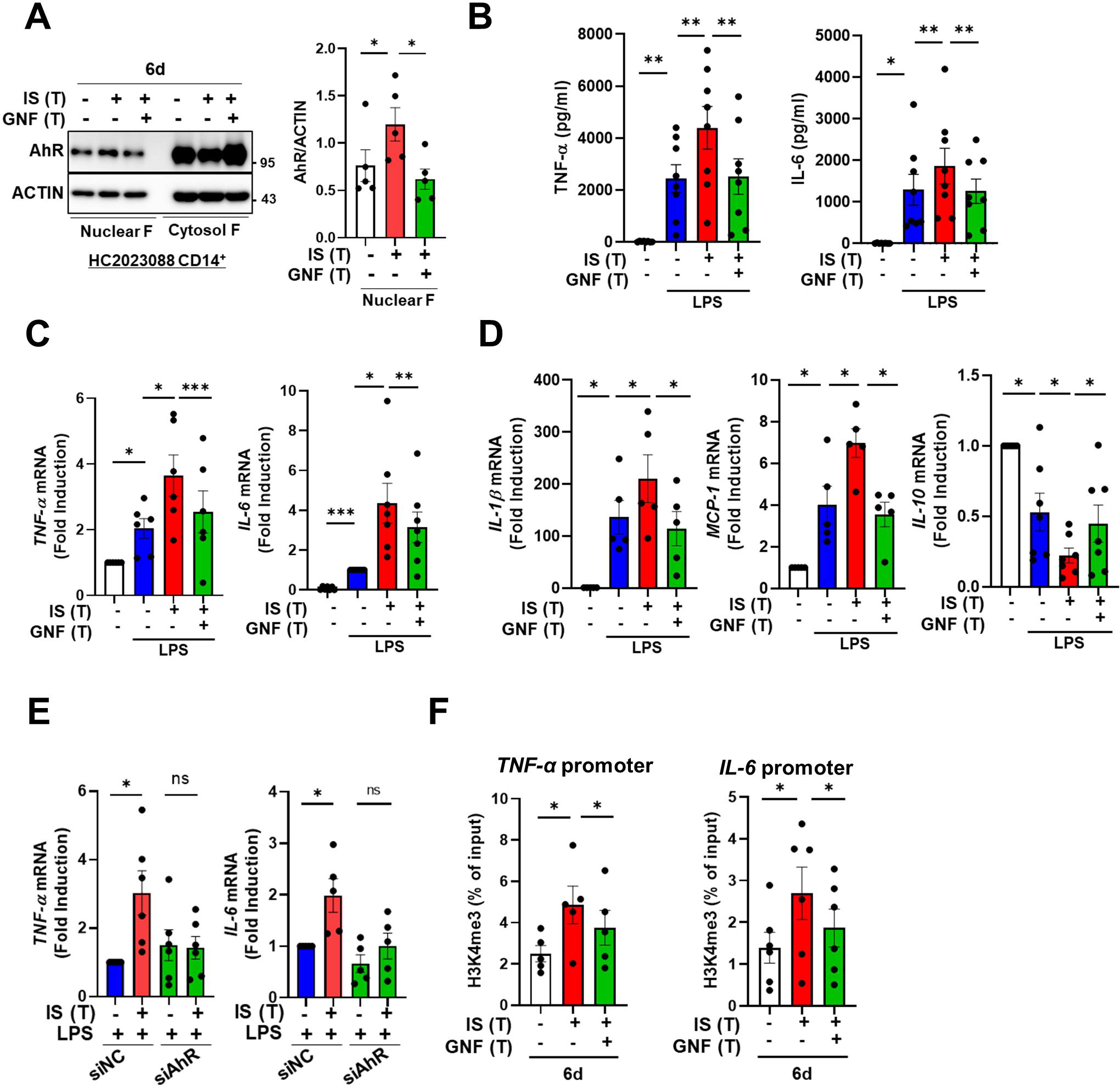
IS-induced trained immunity is regulated by AhR. Monocytes were pretreated with or without GNF351 (AhR antagonist; 10 μM) followed by IS (1,000 μM)-training for 6 days. **A.** On day 6, nuclear and cytosol fraction were prepared and immunoblotted for AhR protein. Band intensity in immunoblots was quantified by densitometry. β-ACTIN was used as a normalization control. **B-D.** On day 6, IS-trained cells with or without GNF351 were restimulated with LPS (10 ng/ml), for 24 hr. TNF-α and IL-6 in supernatants were quantified by ELISA (B). Expression of TNF-α and IL-6 (C) and *IL-1β*, *MCP-1,* and *IL-10* mRNA (D) was analyzed by RT-qPCR. **E.** Monocytes were transfected with siRNA targeting AhR (siAhR) or negative control (siNC) for 1 day, followed by stimulation with IS for 24 hours. After a resting period of 5 days, cells were re-stimulated with LPS for 24 hours. mRNA expression levels of *TNF-*α and *IL-6* were assessed using RT-qPCR. **F.** Enrichment of H3K4me3 on promoters of *TNFA* and *IL6* loci was assessed on day 6 after IS-training. 1% input was used as a normalization control. Bar graphs show the mean ± SEM. * = *p* < 0.05, **= *p* < 0.01, and *** = *p* < 0.001 by two-tailed paired *t*-test.

### AhR-dependent induction of the arachidonic acid pathway is involved in IS-induced trained immunity

To explore which molecular mechanism is involved in the induction of IS-trained immunity, we performed RNA-sequencing (RNA-Seq) on day 6 (prior to restimulation with LPS) in IS-trained human macrophages. A total of 218 differentially expressed genes (DEGs), consisting of 71 upregulated and 147 downregulated genes, were identified in IS-trained macrophages compared to non-trained cells (Fig. 5A and S5A; FC ≥ ± 2, *p* < 0.05). Gene ontology (GO) analysis of these expression data using the Reactome Gene Set is displayed in Figure 5B. IS-trained macrophages had upregulated pathways including those involved in neutrophil degranulation, integrin cell surface interactions, extracellular matrix organization and arachidonic acid metabolism, whereas pathways associated with kinesins, cell cycle, and the Gα(i) signaling pathway were downregulated (Fig. 5B). Considering the key role of the arachidonic acid (AA) pathway in many inflammatory disorders, we decided to focus on this pathway in the induction of trained immunity by IS. Our findings were supported by Gene Set Enrichment Analysis (GSEA) using Molecular Signatures Database (MsigDB), in which genes related to AA metabolism were enriched in IS-trained macrophages compared to non-trained cells and more importantly, upregulated expression of these genes was inhibited by treatment with GNF351 as illustrated by heatmap analysis of major genes related to AA metabolism (Fig. 5C and D and Fig. S5F). Among AA metabolism pathways, the leukotriene metabolic process, but not the cyclooxygenase (COX) pathway, was primarily involved in the induction of IS-mediated trained immunity (Fig. S5B). Confirmatory RT-qPCR analysis on major AA metabolism-related genes was conducted using IS-trained macrophages obtained from independent, HCs (Fig. 5E). The mRNA expression of arachidonate 5-lipoxygenase (*ALOX5*: also known as *5-LOX* or *5-LO*) and arachidonate 5-lipoxygenase activating protein (*ALOX5AP*: also known as *FLAP*), the enzymes catalyzing AA into leukotrienes (a group of pro-inflammatory lipid mediators) (35–37), was higher in IS-trained macrophages than non-trained cells. In addition, the mRNA expression of *LTB4R1* (also known as *BLT1*), a high-affinity receptor for leukotriene B4 (LTB4), was also upregulated. The augmented expression of these AA metabolism-related genes was repressed by GNF351 pretreatment as shown by changes in expression of *CYP1B1*, a typical AhR target gene. Thus, this suggests that the IS-activated AhR pathway is involved in enhanced AA-metabolism in IS-induced trained immunity. Immunoblot analysis validated the upregulation of ALOX5 and ALOX5AP expression in IS-trained immunity, which was subsequently inhibited by GNF351 at the protein level (Fig. 5F). Furthermore, knockdown of AhR suppressed the IS-induced mRNA expression of *ALOX5, ALOX5AP,* and *LTB4R* on day 6 (Fig. 5G). Treatment with FICZ, an AhR agonist known to induce trained immunity (Fig. S4A), elicited increased expression of *ALOX5* and *ALOX5AP*, while treatment with KA, a major protein-bound uremic toxin that does not induce trained immunity (Fig. 1A), did not result in elevation of these genes, thereby implying the significant role of the AhR-AA pathway in IS-trained immunity (Fig. S5C).

**Figure 5.**
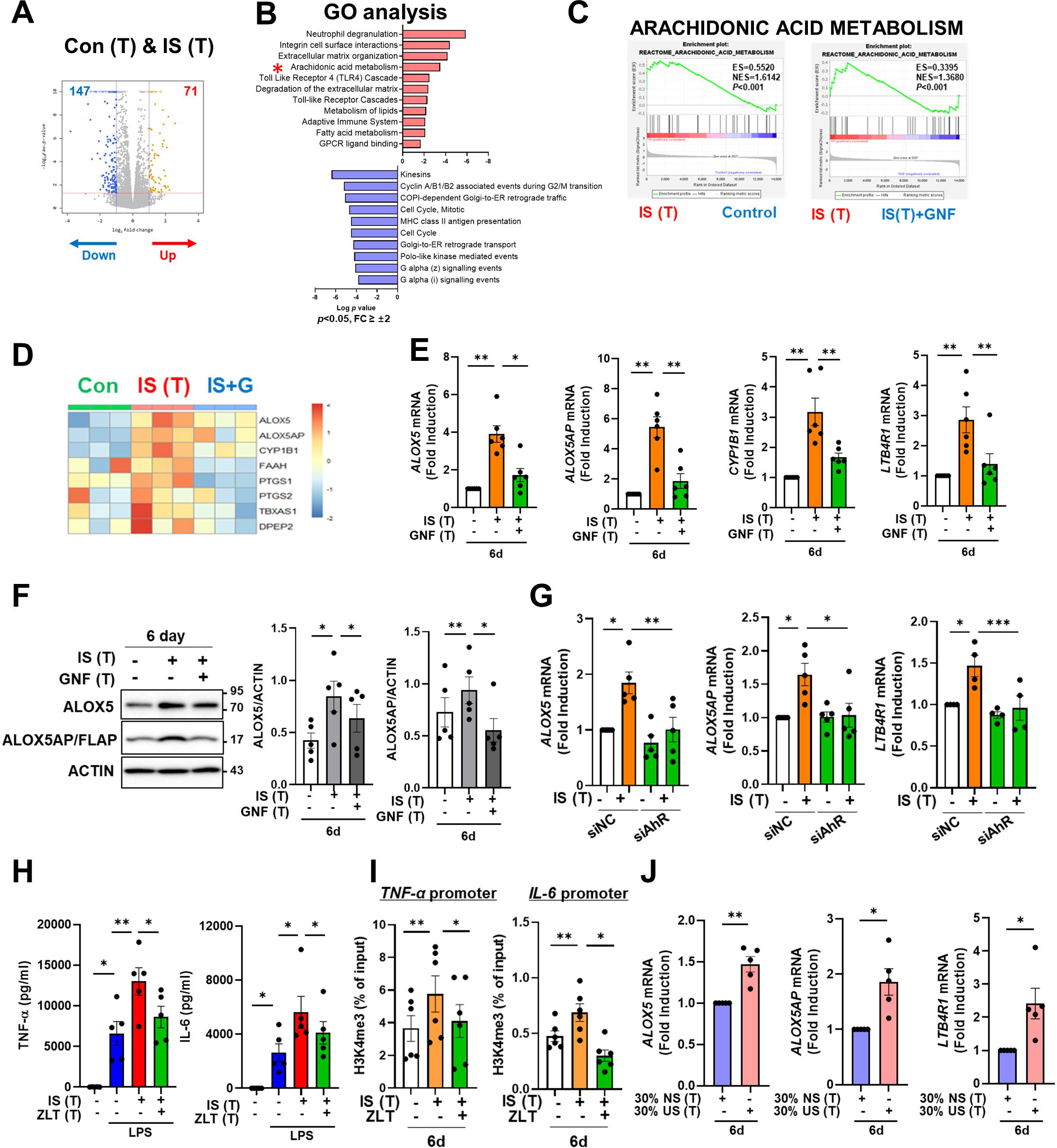
AhR-dependent induction of the arachidonic acid pathway contributes to IS-induced trained immunity. **A.** RNA-Seq analysis was performed on IS (1,000 μM)-trained monocytes. Volcano plots show differentially expressed genes between IS-trained and non-trained macrophages. **B.** Functional annotation of upregulated or downregulated genes (FC > ±2, *p* < 0.05) in IS-trained macrophages analyzed by Gene Ontology (GO) analysis with the Reactome Gene Set. **C-D.** GSEA (C) and heatmap (D) of genes related to the AA metabolism in IS-trained macrophages compared to non-trained cells or compared to IS-trained macrophages with GNF351 (10 μM) treatment were analyzed. **E-F.** On day 6 after IS-training with or without GNF351, expression of *CYP1B1*, *ALOX5*, *ALOX5AP*, and *LTB4R1* mRNAs were quantitated using RT-qPCR (E) and cell lysates were prepared and immunoblotted for ALOX5 and ALOX5AP proteins (F). Band intensity in immunoblots was quantified by densitometry. β-ACTIN was used as a normalization control. **G.** Monocytes were transfected with siRNA targeting AhR (siAhR) or negative control (siNC) for 1 day, followed by stimulation with IS for 24 hours. After a resting period of 5 days, mRNA expression level of each gene was assessed using RT-qPCR. **H.** Monocytes were pretreated with zileuton (ALOX5 inhibitor, 100 μM) and trained with IS for 6 days followed by restimulation with LPS (10 ng/ml) for 24 hr. TNF-α and IL-6 in supernatants were quantified by ELISA. **I.** A ChIP assay was performed in IS-trained macrophages pre-treated with zileuton. 2% input was used as a normalization control. **J.** The pooled normal serum (NS) from health controls (HCs) or uremic serum (US) from patients with ESRD were used to treat monocytes isolated from HCs for 24 hr at 30% (v/v) followed by resting for 5 days. Expression of *ALOX5*, *ALOX5AP*, and *LTB4R1* mRNAs were quantitated using RT-qPCR in trained macrophages with NS or US for 6 days. Bar graphs show the mean ± SEM. * = *p* < 0.05, **= *p* < 0.01, ***= *p*< 0.001 by two-tailed paired *t*-test.

We previously reported alterations in the transcriptome signature of *ex vivo* monocytes of ESRD patients (38). Comparison of the fold changes of RNA-Seq data in the present study and microarray data reported previously (GSE155326) revealed that the expression of *ALOX5* and *LTB4R1* is enhanced in IS-trained macrophages and *ex vivo* monocytes of ESRD patients (Fig. S5D). To further investigate the roles of the AA metabolism pathway in IS-trained immunity, zileuton, an ALOX5 inhibitor, and U75302, a BLT1 receptor inhibitor were used during the induction of trained immunity by IS (Fig. 5H, S5E-F). We found that IS-induced TNF-α and IL-6 production were largely suppressed by both zileuton and U75302 (Fig. 5H, S5F). Additionally, knockdown of ALOX5 inhibited IS-induced expression of TNF-α and IL-6 (Fig. S5H). In further exploration of the effects on epigenetic or metabolic reprogramming *via* the AA pathway, we conducted ChIP-qPCR assays and Western blot analyses following treatment with zileuton. Our results demonstrated that the enrichment of H3K4me3 on *TNFA* and *IL6* promoters in IS-trained macrophages was inhibited by zileuton, although phosphorylation of S6K remained unaffected (Fig. 5I and S5I). Thus, these findings suggest that AA metabolism plays a pivotal role in the induction of IS-trained immunity by serving as a crucial mediator between AhR signaling and epigenetic modification. We next tested whether training with uremic serum leads to increased expression of AA pathway-related genes within 6 days (prior to restimulation with LPS) as found in IS-trained macrophages. The expression of *ALOX5*, *ALOX5AP*, and *LTB4R1* mRNA was augmented by training with pooled uremic sera of ESRD patients compared with HCs (Fig. 5J), and this augmented expression was maintained after re-stimulation with LPS (Fig. S5J).

Histone-modifying enzymes such as lysine demethylase (KDM) and lysine methyltransferase (KMT) are linked to the induction of trained immunity by remodeling the epigenetic status of cells (3, 4). However, RNA-Seq data of IS-trained macrophages showed no obvious change in the expression profile of major histone-modifying enzymes (Fig. S6A). In agreement with this, mRNA expression of major histone modifying enzymes including *KDM5A*, *KDM5B*, *KDM5C*, *SETDB2*, *SETD7*, and *SETD3* were not changed in IS-trained macrophages on day 6 (Fig. S6B) (4, 39–41). Treatment with MTA, a non-selective methyltransferase inhibitor, partially inhibited expression of *ALOX5* and *ALOX5AP* mRNA (Fig. S6C), suggesting limited epigenetic regulation of the AA pathway (Fig. S6C). Ultimately, to validate the association between ChIP-Seq and RNA-Seq data, we employed Spearman’s correlation for comparative analysis and conducted linear regression to ascertain the presence of a consistent global trend in RNA expression. Our findings unveiled a significant positive correlation, underscoring the consistent relationship between H3K4me3 enrichment and gene expression (Fig. S6D).

### IS-induced trained immunity is validated by *ex vivo* and *in vivo models*

Since peripheral monocytes in ESRD patients are chronically exposed to uremic toxins like IS, we examined whether monocytes purified from ESRD patients before hemodialysis exhibit features of IS-trained macrophages. *Ex vivo* monocytes of ESRD patients had a higher production of TNF-α and mRNA expression of *IL-1β* and *MCP-1* after LPS stimulation than those of HCs (Fig. S7A-C). More importantly, monocytes of ESRD patients, which were trained by resting in the culture media for 6 days, significantly augmented production of TNF-α and IL-6 and expression of *IL-1β* mRNA upon LPS stimulation compared to those from HCs (Fig. 6A-C). Consistent with our findings (Fig. 5), the expression of ALOX5 in *ex vivo* monocytes of ESRD patients was significantly increased at the protein level compared with that of age-matched HCs (Fig. 6D-F). Moreover, monocyte-derived macrophages (MDMs) from ESRD patients also had higher expression of ALOX5 compared to MDMs of HCs (Fig. 6G-H), suggesting that IS in serum of ESRD patients contributes to the induction of trained immunity of monocytes/macrophages.

**Figure 6.**
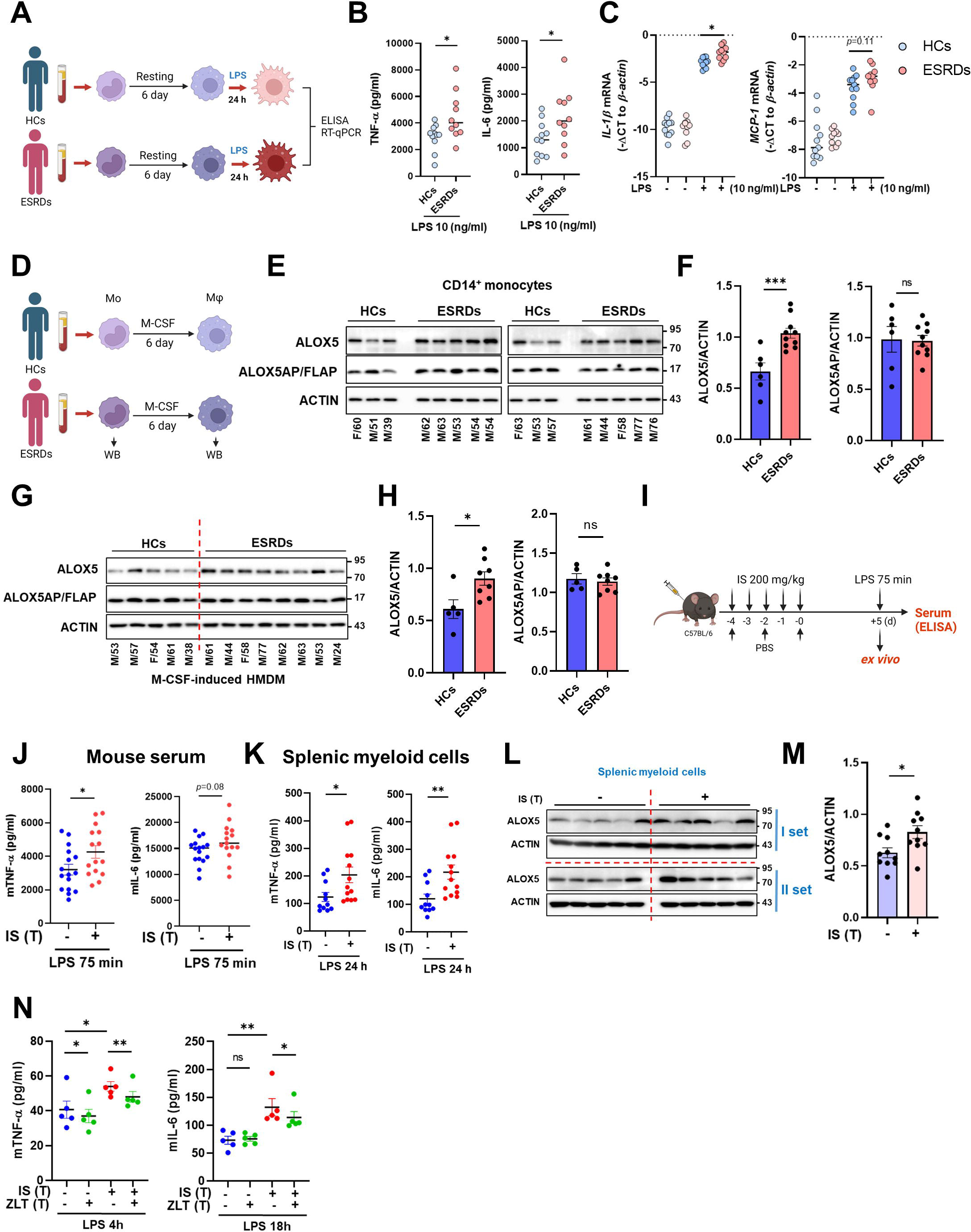
*Ex vivo* and *in vivo* validation of IS-induced trained immunity. **A-C.** CD14^+^ monocytes from ESRD patents and age-matched HCs were rested for 6 days and stimulated by LPS (10 ng/ml) for 24 hrs (A). TNF-α and IL-6 in supernatants were quantified by ELISA (B) and mRNA expression of *IL-1β* and *MCP-1* were quantitated using RT-qPCR (C). **D**-**G.** ALOX5 and ALOX5AP protein levels in monocytes of (E-F) and in M-CSF-derived HMDM (G-H) of ESRD patients and HCs were analyzed by immunoblot analysis. Band intensity in immunoblots was quantified by densitometry. β-ACTIN was used as a normalization control. **I.** C57BL/6 mice were injected daily with 200 mg/kg IS for 5 days and rested for another 5 days prior to LPS (5 mg/kg) treatment. Mice were sacrificed at 75 min post-LPS injection. **J.** TNF-α and IL-6 in serum were quantified by ELISA. **K.** Before LPS injection, IS-trained mice were sacrificed, and spleens were mechanically separated. Isolated splenic myeloid cells were treated *ex vivo* with LPS (10 ng/ml) for 24 hr and TNF-α and IL-6 in supernatants were quantified by ELISA. **L-M.** The level of ALOX5 protein in splenic myeloid cells isolated from IS-trained or control mice was analyzed by western blot. The graph shows the band intensity quantified by the densitometry (M). **N.** Isolated splenic myeloid cells were treated *ex vivo* with LPS (10 ng/ml), along with zileuton (100 µM). The levels of TNF-α and IL-6 in the supernatants were quantified using ELISA. The graphs show the median (B-C) or the mean ± SEM (F-N). *= *p* < 0.05, **= *p* < 0.01, and *** = *p* < 0.001 by unpaired non-parametric *t*-test or by two-tailed paired *t*-test between zileuton treatment group and no-treatment group (N).

To examine the systemic *in vivo* effect of IS-trained immunity, we adopted a murine model in which IS was intraperitoneally injected daily for 5 days, followed by training for another 5 days and then re-stimulation with 5 mg/kg LPS for 75 min (Fig. 6I). The level of TNF-α in serum was increased in IS-trained mice compared to that of control mice (Fig. 6J). To further investigate the impact of IS-training on innate responses, splenic myeloid cells were isolated after 5 days of training (prior to injection with LPS) followed by *in vitro* stimulation with 10 ng/ml LPS for 24 h. The amount of TNF-α and IL-6 in the supernatant was augmented following culture with LPS-stimulated mouse splenic myeloid cells derived from IS-trained mice compared the control condition (Fig. 6K). Additionally, we observed upregulation of ALOX5 expression in *ex vivo* splenic myeloid cells of IS-treated mice compared to control mice (Fig. 6L-M), which was similar to what was observed in monocytes and macrophages from ESRD patients (Fig. 6E-H). Finally, treatment with zileuton, an ALOX5 inhibitor, inhibited the production of TNF-α and IL-6 in *ex vivo* splenic myeloid cells of IS-trained mice (Fig. 6N).

Subsequently, we examined the impact of IS-trained immunity on mouse bone marrow-derived macrophages (BMDM). It was observed that BMDM from IS-trained mice did not exhibit heightened production of TNF-α and IL-6, and the expression level of ALOX5 in bone marrow progenitor cells remained unaltered when compared to non-trained cells, indicating that our acute IS-trained mice did not induce central trained immunity (Fig. S7D-E)(42, 43). Collectively, these findings offer compelling evidence for the involvement of IS and the induction of the AA pathway in the establishment of trained immunity in both monocytes and macrophages, both *ex vivo* and *in vivo*. Together, these data provide evidence for the role of IS and the induction of the AA pathway in the establishment of trained immunity of monocytes and macrophages both *ex vivo* and *in vivo*.

## Discussion

Recent studies have reported that in addition to pathogenic stimuli, endogenous sterile inflammatory insults including ox-LDL, hyperglycemia and uric acid, also trigger trained immunity and contribute to chronic inflammation in cardiovascular diseases and gout (8, 10, 12, 16). In the present study, we provide evidence that IS, a major uremic toxin, provokes trained immunity in human monocytes/macrophage through epigenetic modification, metabolic rewiring, and AhR-dependent induction of the AA pathway, suggesting its important role in inflammatory immune responses in patients with CKD (Fig. 7).

**Figure 7.**
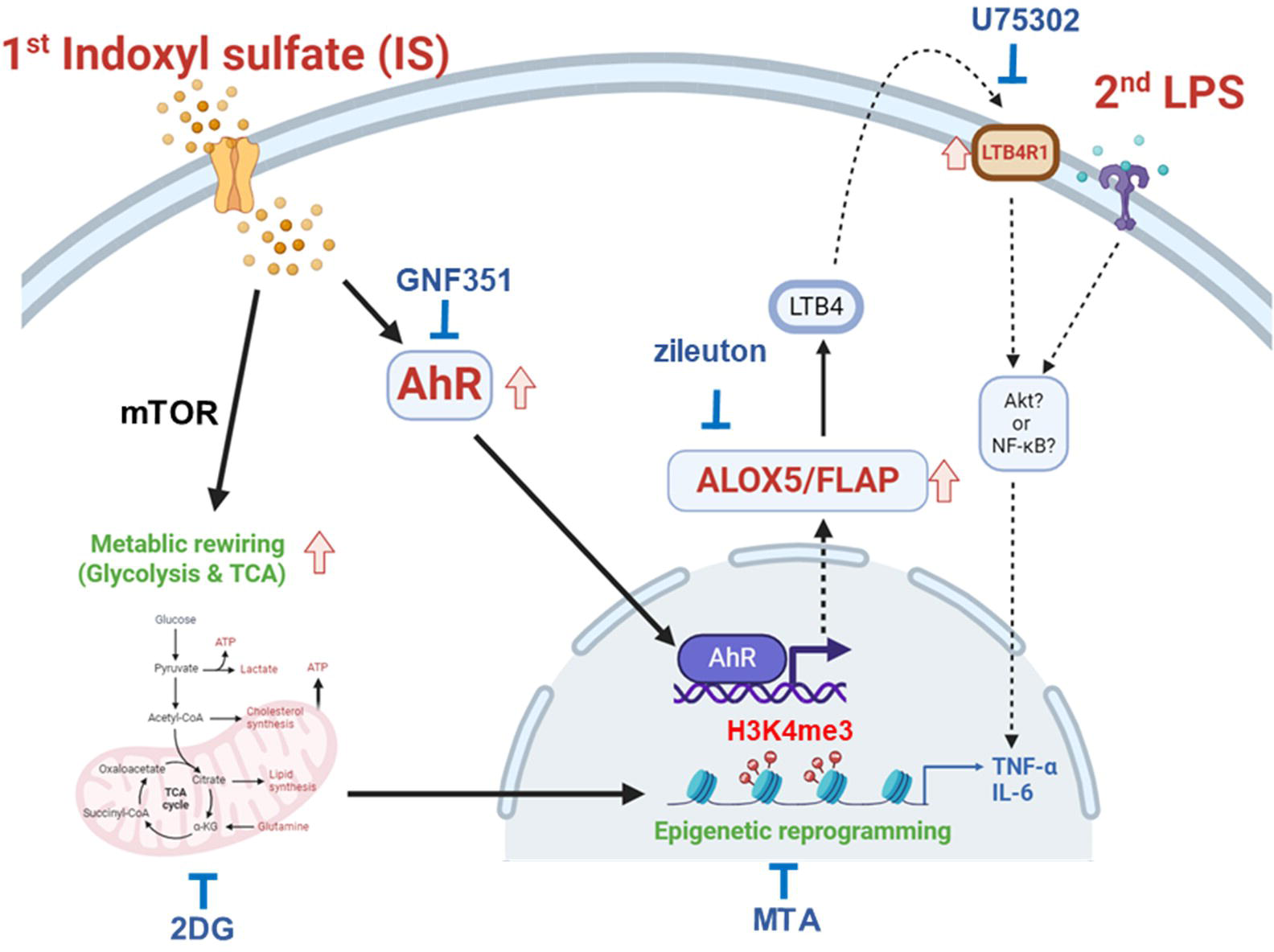
Proposed mechanism of IS-induced trained immunity. IS-induced trained immunity in human monocytes is mediated by epigenetic reprogramming and metabolic rewiring *via* histone modification H3K4m3 and increased glycolysis and mitochondrial respiration, respectively. Direct interaction of uremic toxin IS with the AhR in human monocytes activates AhR signaling pathways that are involved in enhanced expression of the arachidonic acid metabolism-related genes ALOX5, ALOX5AP, and LTB4R1 and augmented production of TNF-α and IL-6 upon stimulation with LPS as secondary stimulus *via* epigenetic regulation. A pivotal role of each pathway or molecule was confirmed by *in vitro* assay with inhibitors including GNF351 (an AhR antagonist), zileuton (an ALOX5 inhibitor), U75302 (a BLT1 receptor inhibitor), 2DG (a glycolysis inhibitor), and MTA (a non-selective methyltransferase inhibitor). Meanwhile, the AhR-independent mechanism contributes to metabolic rewiring, such as increased glycolysis in IS-trained macrophages, which leads to enhanced proinflammatory responses upon secondary stimulation.

CKD is associated with increased risk factors of cardiovascular disease (CVD) including traditional risk factors, such as hypertension, age and dyslipidemia, as well as non-traditional risk factors, such as oxidative stress and inflammation (23, 44). Further, recent cohort studies have shown that CKD is an independent risk factor for CVD (44). Uremia accompanying renal failure causes immune dysfunction, which is closely linked to the pathogenesis of CKD-related CVD (45). Among over 100 uremic toxins identified, IS is a prototypical protein-bound uremic toxin most likely to be participating in progressive pathophysiology of CVD including endothelial dysfunction, vascular calcification, and increased atherosclerosis (21, 26, 46, 47). Mounting evidence suggests that a prolonged hyperactivation of trained immunity is intimately related to the pathogenesis of atherosclerosis, the major contributor to cardiovascular diseases. Oxidized low-density lipoprotein (oxLDL), hyperglycemia, and the Western diet, all known to be associate with the progression of atherosclerosis, have been reported to induce trained immunity through epigenetic reprogramming (8, 10, 12). These findings suggest that IS plays a role as a typical endogenous inflammatory insult in activating monocytes and macrophages and modulating their responses. Given that IS is difficult to clear by hemodialysis, this toxin has a chronic effect on the immune system of patients. Nonetheless, little is known about the effects of IS on trained immunity.

As observed using a common *in vitro* model of trained immunity established by Netea and other groups (Fig. 1A), CD14^+^ monocytes, which are exposed for 24 h to the first insult with IS and rested for 5 days without IS, produced an augmented level of TNF-α and IL-6 and decreased level of IL-10 in response to an unrelated second insult with LPS or Pam3cys, which is a feature typical of trained immunity of monocytes (Fig. 1B-E). In contrast to TNF-α and IL-6, major proinflammatory cytokines, IL-10 exerts potent deactivating effects on macrophages and T cells, influencing various cellular processes in inflammatory diseases (48, 49). Additionally, it is noteworthy that IL-10-deficient macrophages exhibit an augmentation in the proinflammatory cytokine TNF-α (50, 51). Therefore, the reduced gene expression of IL-10 by IS-trained monocytes may contribute to the heightened expression of proinflammatory cytokines. Mechanistic studies have demonstrated that the induction of trained immunity is coordinated through the interplay of epigenetic modifications and metabolic rewiring, which is broadly characterized as prolonged changes in transcription programs and cell physiology that do not involve permanent genetic changes, such as the mutations and recombination events crucial for adaptive immunity (2). In the present study, ChIP-Seq and real-time metabolic analysis show that the induction of IS-trained immunity in human monocytes is attributable to epigenetic modification and metabolic rewiring (Fig. 2 and 3). Consistent with previous findings (5, 12), trimethylation of histones at H3K4 on *TNFA* and *IL6* promoters was increased in IS-trained macrophages and maintained even after secondary stimulation with LPS (Fig. 3B and Fig. S3B). Furthermore, the production of TNF-α and IL-6 upon LPS stimulation was completely inhibited by pretreatment with 5’-methylthioadenosine (MTA), a methyltransferase inhibitor (Fig. 3C), demonstrating that IS-induced trained immunity is associated with epigenetic modification. Moreover, H3K4me3-ChIP-Seq data showed that IS-induced trained immunity accompanied by epigenetic reprogramming and H3K4me3 was enriched in genes related to activation of the innate immune responses as illustrated by gene ontology (GO) analysis (Fig. 3E-G).

We identified AhR as a critical mediator of IS-trained immunity in human monocytes (Fig. 4). AhR is a ligand-activated nuclear transcription factor (TF), which is activated by several exogenous compounds, such as benzo[a]pyrene environmental pollutants and 2,3,7,8-tetrachloro-dibenzo-*p*-dioxin (TCDD), as well as by multiple endogenous ligands including tryptophan and indole metabolites(52). AhR plays a multifaceted role in modulating cellular mechanisms such as inflammation, cell growth, and antioxidant responses (53). AhR is expressed by various immune cells, and its signaling exerts integrative effects on the cellular environment and metabolism of the immune responses (54). However, little is known about the role of AhR in the induction of trained immunity.

In the present study, we show that IS-trained immune responses, characterized by the expression of proinflammatory cytokines and chemokines, were attenuated by GNF351, an AhR antagonist, or through the knockdown of AhR using siRNA (Fig. 4B-E). This inhibition was accompanied by repression of enriched H3K4me3 on *TNFA* and *IL6* promoters in IS-trained macrophages (Fig. 4F), indicating an AhR-dependent mechanism. However, increased glycolysis and mitochondria respiration in IS-trained macrophages were not suppressed by the blockade of AhR activation with GNF351, suggesting the AhR activation is not directly involved in metabolic rewiring in IS-trained immunity (Fig. S4B-D).

Metabolic rewiring, especially upregulation of aerobic glycolysis, is known as a major mechanism underlying the induction of trained immunity *via* regulation of epigenetic modification by metabolites, such as mevalonate and fumarate generated from this metabolic rewiring (55). Furthermore, in addition to establishing the bidirectional link between heightened glycolysis and activating histone modifications, as evidenced by the suppression of histone modification by 2DG, a glycolysis inhibitor, in IS-induced immunity (Fig. 3D), recent studies have demonstrated that immune priming long non-coding RNAs (IPLs) also induce epigenetic reprogramming by influencing 3D nuclear architecture(3, 56) and that RUNX1, a transcription factor, contributes to the induction of trained immunity by overexpression of RUNX1 target genes (10), suggesting that a variety of mechanism is involved in epigenetic modification in trained immunity.

Although inducers of trained immunity, such as β-glucan, BCG, uric acid, and oxLDL, initiate intracellular signaling and metabolic pathways, each *via* different receptors, the most common pathway is the Akt/mTOR/HIF1α-dependent induction of aerobic glycolysis (3, 15, 55). Our data also revealed that training with IS led to enhanced glycolysis, which is critical for the production of TNF-α and IL-6 upon LPS stimulation as confirmed by experiments with 2DG, a glycolysis inhibitor (Fig. 2E). Recent studies have shown that IS activates mTORC1 in a variety of cells such as epithelial cells, fibroblasts, and THP-1 cells mainly *via* the Organic Anion Transporters (OAT)/NADPH oxidase/ROS pathway, but not the AhR pathway (57). Our findings suggest that IS-trained macrophages acquire the characteristics of trained immunity by AhR-dependent and -independent mechanisms and enhances proinflammatory responses upon secondary stimulation. Thus, addressing the mechanism underlying AhR-independent metabolic rewiring in IS-trained macrophages will require further investigations.

A finding of particular interest in our study is that the induction of IS-induced trained immunity is dependent on the AhR-ALOX5/ALOX5AP axis, as depicted by RNA-Seq analysis and confirmatory *in vitro* analysis of mRNA and protein expression (Fig. 5 and Fig. S5). ALOX5 and ALOX5AP are major rate-limiting enzymes associated with the arachidonic acid (AA) pathway, involved in the production of leukotrienes, proinflammatory lipid mediators derived from AA (36). Among leukotrienes, leukotriene B4 (LTB4), an extremely potent inflammatory mediator, binds to G protein-coupled protein, LTB4R, and enhances inflammatory responses by increased phagocytosis and activation of the signaling pathway for the production of cytokines (35, 37). Mechanistically, the LTB4-LTB4R signaling pathway induces the PI3K/Akt or NF-κB pathways (58, 59). AhR-dependent LTB4 production through enhanced ALOX5 expression in hepatocytes reportedly induces hepatotoxicity *via* neutrophil infiltration (60). The increase in ALOX5 activity and LTB4 expression has been reported in patients with ESRD (61–63) and ALOX5 mediates mitochondrial damage and apoptosis in mononuclear cells of ESRD patients (61). Thus, antagonists of ALOX5/ALOX5AP have been used for treatment in CKD (63). Consistent with these findings, peripheral monocytes derived from ESRD patients in our cohort have increased expression of ALOX5 and this increase was maintained after differentiation into macrophages with M-CSF (Fig. 6E-H). Considering that pretreatment with the uremic serum of ESRD patients elicits a change in gene expression and cytokine production as observed in IS-trained macrophages, it is likely that increased ALOX5 in monocytes and macrophages of ESRD patients is mediated by IS in uremic serum. Our previous studies have shown that IS is an important uremic toxin in the serum of ESRD patients that elicits proinflammatory responses of monocytes (21). Furthermore, the AhR-ALOX5-LTB4R1 pathway is involved in IS-induced trained immunity of patients with ESRD.

Recent studies have demonstrated that trained immunity of macrophages is implicated in the pathogenesis of CVD, such as atherosclerosis (10, 12, 13, 15). Considering a higher plasma level of IS in ESRD patients even after hemodialysis (22, 23), this is potentially difficult to reconcile with trained immunity in which there typically is a short exposure to the training stimulus followed by a period of rest (2, 3). However, when ESRD monocytes exposed to the IS in the circulation enter atherosclerotic plaques and differentiate into macrophages, they are no longer exposed to IS, because this is protein-bound and hence not expected to the present in the plaque microenvironment.

Murine models have been widely used to investigate the specific contribution of inducers of trained immunity and the underlying mechanisms to a long-term functional modification of innate cells *in vivo* (7, 41, 64). Mice trained with β-glucan enhance the production of proinflammatory cytokines such as TNF-α, IL-6, and IL-1β by monocytes and macrophages in response to secondary microbial stimuli and subsequently obtain increased protection against various microbial infections (15, 55). Our *in vivo* and *ex vivo* mouse experiments for trained immunity demonstrate that IS-trained immunity has biological relevance (Fig. 6I). Previous studies have shown that intraperitoneal injection of exogenous IS into wild-type C57BL/6 mice leads to an increase of IS in the plasma until 3∼6 hr post-injection despite its rapid excretion by the kidney. In this period, the expression of pro-inflammatory, pro-oxidant, and pro-apoptotic genes in peritoneal macrophages was upregulated (57, 65). Moreover, intraperitoneal injection of IS daily for 3 days elevates IS in the plasma of mice and activates the mTORC1 signaling pathway in the mouse kidney (57). This suggests that the IS-injected mouse model is suitable for investigating the mechanisms underlying IS-mediated immune responses. Our data show increased TNF-α in serum after injection of LPS (Fig. 6J). Moreover, ALOX5 protein expression was increased in splenic myeloid cells derived from IS-trained mice and *ex vivo* stimulation with LPS for 24 hr induced TNF-α and IL-6 expression in these splenic myeloid cells (Fig. 6K-M). Thus, this suggests a systemic induction of IS-trained immunity in the mouse model.

In conclusion, the current study provides new insight into the role of IS as an inducer of trained immunity as well as the underlying mechanisms in human monocytes/macrophages by investigating the effect of IS *in vivo* and *in vitro* using experimental models of trained immunity. Here, we demonstrate that IS, a major uremic toxin, induces trained immunity characterized by the increased proinflammatory TNF-α and IL-6 in human monocytes following secondary stimulation through epigenetic modification and metabolic rewiring. IS-mediated activation of AhR is involved in the induction of trained immunity through enhanced expression of arachidonic acid (AA) metabolism-related genes such as ALOX5 and ALOX5AP. Monocytes from patients with ESRD exhibit increased expression of ALOX5 and after 6-day resting, they exhibit enhanced TNF-α and IL-6 production to LPS. Furthermore, the uremic serum of ESDR patients causes HC-derived monocytes to increase the production of TNF-α and IL-6 upon LPS re-stimulation, implying IS-mediates trained immunity in patients. Supporting our *in vitro* findings, mice trained with IS and their splenic myeloid cells had increased production of TNF-α after *in vivo* and *ex vivo* LPS stimulation. These results suggest that IS plays an important role in the induction of trained immunity, which is critical in inflammatory immune responses in patients with CKD and thus, it holds potential as a therapeutic target.

## Methods

### Human monocyte preparation and culture

Peripheral blood mononuclear cells (PBMCs) were isolated from peripheral blood by density gradient centrifugation (Bicoll-Separating Solution; BIOCHROM Inc, Cambridge, UK). Monocytes were positively purified from PBMCs with anti-CD14 magnetic beads (Miltenyi Biotec Inc, Auburn, CA). For *in vitro* trained immunity experiments, purified monocytes were treated with IS for 24 h, followed by washing with pre-warmed PBS and incubation for another 5 days in RPMI medium supplemented with 10% human AB serum (HS, Sigma-Aldrich, St. Louis, MO), 100□U/ml penicillin, and 100□μg/ml streptomycin (Gibco, Grand Island, NY). On day 6, cells were re-stimulated with LPS or Pam3cys for 24 hr, and the supernatants and their lysates were collected and stored at -80□ until use. In some experiments, chemical inhibitors were used for a 1 hr pre-treatment at the indicated concentrations prior to the treatment with IS. To test the effect of uremic serum on the induction of trained immunity, CD14^+^ monocytes purified from HC donors were seeded into 48-well plates and incubated for 24 hr at 30% (v/v) with the pooled uremic sera (US) from ESRD patients or the pooled normal sera (NS) from HCs, followed by washing with pre-warmed PBS and incubation for another 5 days. On day 6, cells were re-stimulated with LPS for 24 hr. To examine whether monocytes from ESRD patients inhibit features of IS-trained immunity, CD14^+^ monocytes were purified from ESRD patients and age-matched HCs, followed by stimulation with LPS for 24 hrs. In addition, purified monocytes were seeded and rested for 6 days in RPMI medium supplemented with 10% human AB serum, 100□U/ml penicillin, and 100□μg/ml streptomycin to induce trained immunity. On day 6, the cells were stimulated with LPS for 24 hr. In some experiments, monocyte-derived macrophages (MDMs) were differentiated from purified CD14^+^ monocytes from ESRD patients or age-matched HCs in RPMI 1640 medium supplemented with 10% fetal bovine serum (FBS, BioWest, Nuailĺe, France), 50 ng/ml recombinant human M-CSF (PeproTech, Rocky Hill, NJ, USA), 100 U/ml penicillin, and 100 mg/ml streptomycin. On day 6, MDMs were used for immunoblot analysis.

### Chemicals and antibodies

Indoxyl sulfate (IS) potassium salt, GNF351, 5’-methylthioadenosine (MTA), 2-deoxy d-glucose (2DG), and zileuton were purchased from Sigma-Aldrich (Burlington, MA, USA). Lipopolysaccharides (LPS) from E. coli 0111: B4 were purchased from InvivoGen (San Diego, CA, USA) for *in vitro* experiments and Sigma-Aldrich for *in vivo* experiments. U-75302 was obtained from Cayman Chemical (Ann Arbor, Michigan, USA). Anti-AhR and anti-5-loxygenase (5-LOX) antibodies (Ab) for immunoblot assay and anti-trimethyl H3K4 (H3K4me3) Ab for chromatin immunoprecipitation (ChIP) were purchased from Cell Signaling Technology (Danvers, MA, USA). Anti-ALOX5AP (FLAP) antibody was obtained from Abcam Inc. (Cambridge, UK).

### Enzyme-linked immunosorbent assay (ELISA)

The amounts of TNF-α and IL-6 in culture supernatants of LPS or Pam3cys-re-stimulated IS-trained macrophages were quantified using commercial human ELISA kits (Thermo Fisher Scientific, Waltham, MA, USA). Optical density was measured using the Infinite M200 (Tecan, Männedorf, Switzerland).

### Quantitative RT-PCR

Total RNA was prepared using RNA purification kit (Macherey-Nagel GmbH & Co. KG, Germany), followed by cDNA synthesis (Bio-line, London, UK), and then real-time quantitative RT-PCR was performed with the CFX system (Bio-Rad, Hercules, CA) using the SensiFAST SYBR® Lo-ROX (Bio-line, London, UK). Sequences of primers used in this investigation are shown in Table S3. Normalization of gene expression levels against the expression of ACTINB using the comparative CT method (ΔΔCT) was used for quantification of gene expression.

### ChIP-qPCR and ChIP-Seq

Cells were washed with Dulbecco’s PBS and crosslinked for 5 min with 1% formaldehyde at room temperature (RT), followed by quenching with 100 mM glycine for 5 min. Cells were harvested with lysis buffer (50 mM HEPES, pH7.5, 140 mM NaCl, 1 mM EDTA, 10% glycerol, 0.5% NP-40 and 0.25% Triton X-100) with protease inhibitors on ice for 10 min and were then washed with washing buffer (10 mM Tris-HCl, pH7.0, 200 mM NaCl, 1 mM EDTA, and 0.5 mM EGTA) for 10 min. The lysates were resuspended and sonicated in sonication buffer (10 mM Tris-HCl, pH8.0, 100 mM NaCl, 1 mM EDTA, 0.5 mM EGTA, 0.1% sodium deoxylcholated and 0.5% N-laurolsarcosine) using a Bioruptor® (diagenode, Denville, NJ) with 30s on and 30s off on a high-power output for 25 cycles. After sonication, samples were centrifuged at 12,000 rpm for 10 min at 4 □ and 1% sonicated cell extracts were saved as input. Cell extracts were incubated with protein A agarose loaded with the H3K4me3 Ab overnight at 4 □, and then Ab-bound agarose beads were washed twice with sonication buffer, once with sonication buffer with 500 mM NaCl, once with LiCl wash buffer (10 mM Tris-HCl, pH8.0. 1 mM EDTA, 250 mM LiCl and 1% NP-40), and once with TE with 50 mM NaCl. After washing, DNA was eluted in freshly prepared elution buffer (1% SDS and 0.1 M NaHCO_3_). Cross-links were reversed by overnight incubation at 65□ with RNase A, followed by incubation with proteinase K for 1 hr at 60□. DNA was purified with NucleoSpin™ gDNA Clean-up Kit (Macherey-Nagel GmbH & Co. KG, Germany). For ChIP-qPCR assays, immunoprecipitated DNA was analyzed by quantitative real-time PCR and results were normalized against input DNA. The sequences of primers used for ChIP-qPCR are shown in Table S4 (5, 66).

For ChIP-seq experiments, purified DNA were prepared for DNA libraries using TruSeq DNA Sample Prep Kit according to Library Protocol TruSeq ChIP Sample Preparation Guide 15023092 Rev. B. Next, illumina sequencing were performed using NovaSeq 6000 S4 Reagent Kit according to sequencing protocol of NovaSeq 6000 System User Guide Document # 1000000019358 v02. Sequenced reads were trimmed using Trimmomatic software. Fragments were aligned to hg19 using Bowtie2 software. Aligned fragments of H3K4me3-ChIP samples were concatenated into a single file to generate consistent peak ranges between samples using the makeTagDirectory function of Homer Suite. For each sample, regions of H3K4me3 enrichment compared to the input sample were collected using callpeaks function in MACS3 software. H3K4me3-rich regions from the same group of different donors were compared to peaks in linked samples using the findoverlap function of the GenomicRange R-package, and 11,123 peaks were collected for further analysis. For quantitative comparisons between IC-trained groups and controls, the number of fragments of each peak in BEDPE was collected using the coverage function of the BEDtools software. Then, the number of fragments in the peak was normalized to CPM and significance was compared using edgeR R-package. Finally, we selected 7,136 peaks with at least 15 CPM from the larger average group to exclude lowly H3K4me3 enriched peaks.

Enriched peaks were selected base on a *p*-value of 0.05 or less and log2 fold change of > 1.3. The selected enriched peaks were used for Go pathway analysis. Pathway analysis was conducted using Metascape web-based platform (67) and significant pathways were identified on the basis of Go biological process and Reactome gene sets. Significant pathways were selected with *p* < 0.05 and enrichment score (ES) > 1.5.

### RNA-sequencing (RNA-Seq) and analysis

After RNA extraction, libraries for sequencing were prepared using the TruSeq Stranded mRNA LT Sample Prep Kit and sequencing were performed using NovaSeq 6000 System User Guide Document # 1000000019358 (Illumina). To analyze RNA-Seq data, trimmed reads were aligned to the human GRCh37 (NCBI_105.20190906). Gene expression profiling was performed using StringTie and then read count and FPKM (Fragment per Kilobase of transcript per Million mapped reads) were acquired. Differentially expressed genes (DEGs) were selected based on *p*-value of 0.05 or less. Selected data were applied to hierarchical cluster analysis to display basal and luminal differences and were further filtered according to gene expression levels with a log2 fold change of < -2 and > 2. DEGs were visualized using the R (ver. 4.1.1) and pheatmap package (ver. 1.0.8). For Gene Set Enrichment Analysis (GSEA), samples were categorized into distinct groups based on treatment conditions. All transcripts within annotated genes (∼14,404 features in total) regarding expression values were uploaded to locally-installed GSEA software (ver. 4.2.3) (68). The analysis was performed by comparing the ‘IS (T)’ group against the ‘Control’ group to identify differentially enriched gene sets within the Reactome pathway database, particularly focusing on the arachidonic acid metabolism pathway. A similar comparison was made between the ‘IS (T)’ and ‘IS (T)+GNF’ groups to assess the effect of the GNF inhibitor. Outputs were filtered based on a nominal *p* < 0.05 and a normalized enrichment score (NES) > 1.3 to determine statistical significance. These thresholds were applied to ensure the robustness of the findings in the context of multiple hypothesis testing. The enrichment plots were generated to visualize the distribution of the gene sets and their enrichment scores. Lastly, pathway analyses were further substantiated using the Metascape web-based platform(67), with significant pathways identified using differentially expressed genes (DEGs) and selected based on a *p* < 0.05 and an enrichment score (ES) > 1.5.

### Metabolic analysis

To profile the metabolic state of the cells, CD14^+^ monocytes were seeded onto XFe24 cell culture plates (Seahorse Bioscience, Lexington, MA) with RPMI medium with 10% HS, followed by the induction of trained immunity for 6 days as described in Fig. 1A. Metabolic analysis on IS-trained macrophages was performed according to the manufacturer’s instructions. For the glycolysis stress test, culture media was replaced with Seahorse XF Base media supplemented with 2 mM L-glutamine (pH7.4) and incubated for 1 hr in the non-CO2 incubator. Glucose (10 mM), oligomycin (2 μM), and 2-DG (50 mM, all from Sigma-Aldrich) were sequentially used to treat cells during real-time measurements of extracellular acidification rate (ECAR) using Seahorse XFe24 Analyzer (Seahorse Bioscience). For the mito stress test, cells were incubated with Seahorse XF Base media supplemented with 1 mM pyruvate, 2 mM L-glutamine, and 10 mM glucose (pH7.4) for 1 hr in the non-CO2 incubator. Oligomycin (1.5 μM), FCCP (2 μM), and rotenone/antimycin A (0.5 μM, all from Sigma-Aldrich) were sequentially used to treat cells during real-time measurements of oxygen consumption rate (OCR) using the Seahorse XFe24 Analyzer. Parameters of glycolysis stress test and mito stress test were calculated using Seahorse XF glycolysis or the mito stress test report generator program that was provided by the manufacturer.

### Immunoblot analysis

Total proteins were prepared using radioimmunoprecipitation assay (RIPA) buffer (150 mM NaCl, 10 mM Na_2_HPO_4_, 0.5% sodium deoxycholate, 1% NP-40) containing a protease and phosphatase inhibitor cocktail (Thermo Fisher Scientific, Waltham, MA, USA). Cell lysates were separated on an 8-12% SDS-polyacrylamide gel and blotted onto a polyvinylidene difluoride (PVDF) membrane (Bio-Rad, Hercules, CA, USA), The membrane was incubated overnight at 4°C with primary Abs, such as anti-AhR, anti-ALOX5, and anti-ALOX5AP/FLAP, followed by incubation with peroxidase-conjugated secondary Abs for 1 h. The membranes were developed using the enhanced chemiluminescence (ECL) system.

### WST (Water Soluble Tetrazolium Salt) assay

To test cell viability, IS-trained macrophages were re-stimulated with LPS for 24 hr. Culture media was changed with serum-free RPMI medium and the WST reagent (EZ-CYTOX, DoGenBio, Seoul, Korea) followed by incubation for 1-2 h. Measurement of the optical density value (450 nm) was performed by Infinite M200 (Tecan).

#### Mouse *in vivo* studies

For *in vivo* experiments, C57BL/6 mice (7-8 weeks) were injected intraperitoneally with 200 mg/kg IS in 100 μl PBS daily for 5 days. Another five days after IS injection, 5 mg/kg LPS (Sigma-Aldrich) were injected intraperitoneally 75 min prior to sacrifice. Whole blood was incubated at RT for 30 min and centrifuged at 3,000 g for 10 min at 4□ to collect mouse serum. The amount of TNF-α and IL-6 in serum was quantified using commercial mouse ELISA kits (Thermo Fisher Scientific). For *ex vivo* experiments using splenic myeloid cells, IS-trained mice were sacrificed and their spleens were aseptically collected. Single-cell splenic suspensions were prepared in PBS after passage through a 40 mm cell strainer. Splenocytes were seeded at 1 x 10^7^ cells/well in 12-well plates. After incubation for 1 h, adherent cells were harvested for immunoblot analysis or stimulated with 10 ng/ml LPS for 24 hr. The amount of TNF-α and IL-6 in culture supernatants was quantified using commercial mouse ELISA kits (Thermo Fisher Scientific).

### Statistical Analysis

A two-tailed paired or unpaired non-parametric t-test was performed to analyze data using Prism 8 (GraphPad Software, La Jolla, CA, USA) and Microsoft Excel 2013. *P* values of less than 0.05 were considered statistically significant.

### Study approval

Study protocols were reviewed and approved by the IRB (institutional review board) of Seoul National University Hospital and Severance Hospital. Peripheral blood of ESRD patients and healthy controls (HCs) was drawn after obtaining written, informed consent. The methods were performed in accordance with the approved guidelines.

## Author contributions

H. Y. Kim: conceived of the study, participated in its design and coordination, performed most of the experiments, data collection, and analysis, wrote manuscript and provided financial support. Y. J. Kang, D. H. Kim, J. Jang, S. J. Lee, and G. Kim: performed the experiments and data analysis. H. B. Koh and Y. E. Ko: performed patient sample collection, the experiments, and data analysis. H. M. Shin: participated in its design and performed data analysis. H. Lee and T.-H. Yoo: conceived of the study, participated in its design, collected patient samples and clinical information, and performed data analysis. W-W.L.: conceived of the study, participated in its design and coordination, performed data analysis and writing of manuscript, and has full access to all the data in this study and financial support. All authors have read and approved the final manuscript.

## Acknowledgments

The authors thank the Core Lab, Clinical Trials Center, Seoul National University Hospital for drawing blood. This work was supported in part by a grant (Grant no: 2022R1A4A1033767 and 2022R1A2C3011243 to W-W. Lee) from the National Research Foundation of Korea (NRF) funded by Ministry of Science and ICT (MSIT) and by a grant (Grant no: RS-2023-00238632 to H.Y.K.) of Basic Science Research Program through NRF funded by the Ministry of Education, Republic of Korea.

