## Supplemental figure 1-7, Supplemental table 1-4 for "Uremic toxin indoxyl sulfate induces trained immunity *via* the AhR-dependent arachidonic acid pathway in end-stage renal disease (ESRD)"

**This file includes:**

Figs. S1 to S7

Tables S1 to S4

Fig. S1.

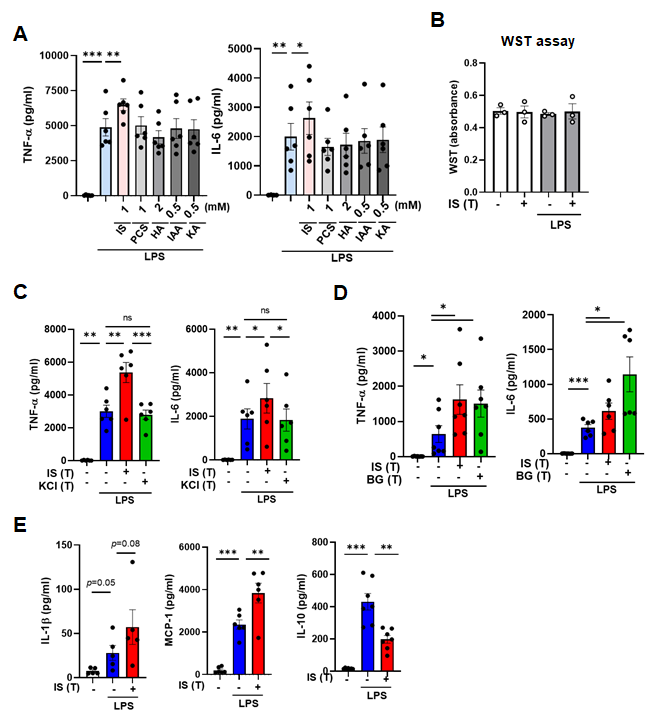

**Supplementary Figure 1.** **IS induces trained immunity in human monocytes. A.** IS (1 mM), PCS (1 mM), Hippuric acid (HA, 2 mM), indole 3-acetic acid (IAA, 0.5 mM), or Kynurenic acid (KA, 0.5 mM) were used to treat cells for 24 hr followed by resting for 5 days. Trained macrophages were restimulated with LPS at 10 ng/ml for 24 hr as described in Figure 1A. TNF-α and IL-6 proteins levels were quantified by ELISA. **B.** Cell death of IS-trained macrophages was analyzed using WST assay. **C.** Monocytes were pretreated with IS (1 mM) or KCl (1 mM) as a vehicle for 24 hr, followed by training for 5 days. Cells were restimulated with 10 ng/ml LPS for 24 hr. TNF-α and IL-6 in supernatants were quantified by ELISA. **D.** β-glucan (10 μM) or IS was pretreated for 24 hr, followed by resting for another 5 days. On day 6, cells were restimulated with 10 ng/ml LPS for 24 hr. TNF-α and IL-6 in supernatants were quantified by ELISA. **E.** Trained macrophages were restimulated with LPS at 10 ng/ml for 24 hr. IL-1β, MCP-1, and IL-10 proteins levels were quantified by ELISA. Bar graphs show the mean ± SEM. *= *p* <0.05, ** = *p* < 0.01, and *** = *p* <0.001 by two-tailed paired *t*-test.

Fig. S2.

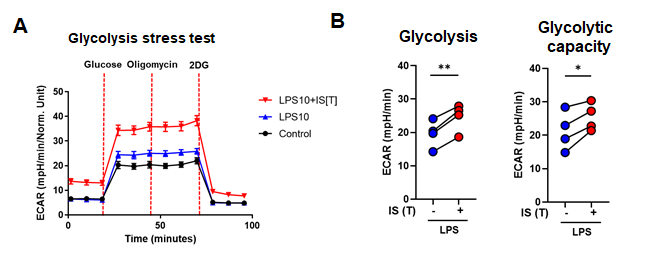

Supplementary Figure 2. IS-induced trained immunity is linked to metabolic rewiring. Related to Figure 3. Glycolysis stress test was conducted using the Seahorse XF-analyzer with IS (1,000 μM)-trained macrophages (n = 4) restimulated with LPS (10 ng/ml). A. ECAR (extracellular acidification rate) levels were measured after sequential treatment with glucose, oligomycin, and 2-DG. B. Cellular glycolysis and glycolytic capacity were analyzed. *= *p* < 0.05 and **= *p* < 0.01 by two-tailed paired *t*-test.

Fig. S3.

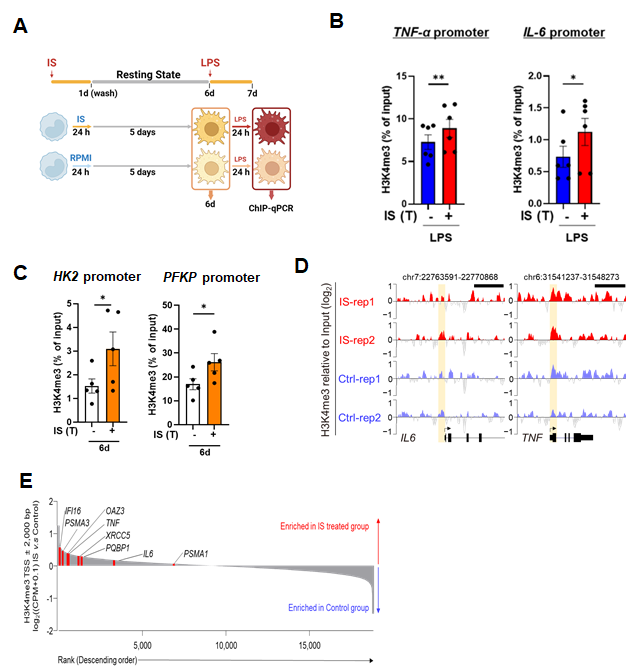

**Supplementary Figure 3. IS-induced trained immunity is associated with epigenetic modification in human innate immune cells.** **A.** Experimental scheme of ChIP-qPCR for IS (1,000 μM)-trained macrophages. **B.** IS-trained macrophages were restimulated with LPS (10 ng/ml) for 24 hr and then cells were fixed with 1% formaldehyde, lysed, and sonicated. ChIP assay was performed using anti-H3K4me3 antibody and enrichment of H3K4me3 at the promoter site of TNFA and IL6 locus was quantified by qPCR. 2% input was used as a normalization control. **C.** On day 6 after IS-training, ChIP assay was performed using anti-H3K4me3 antibody and enrichment of H3K4me3 at the promoter site of *HK2* and *PFKP* loci was quantified by qPCR. 1% input was used as a normalization control. **D-E.** A whole-genome assessment of the histone marker H3K4me3 was analyzed by ChIP-sequencing (ChIP-Seq) in IS-trained cells on day 6. H3K4me3 peak of promoter region on *TNFA* and *IL6* (D). The differences in H3K4me3 enrichment patterns between control group and IS-training group (E). Bar graphs show the mean ± SEM. *= *p* < 0.05 by two-tailed paired *t*-test.

Fig. S4.

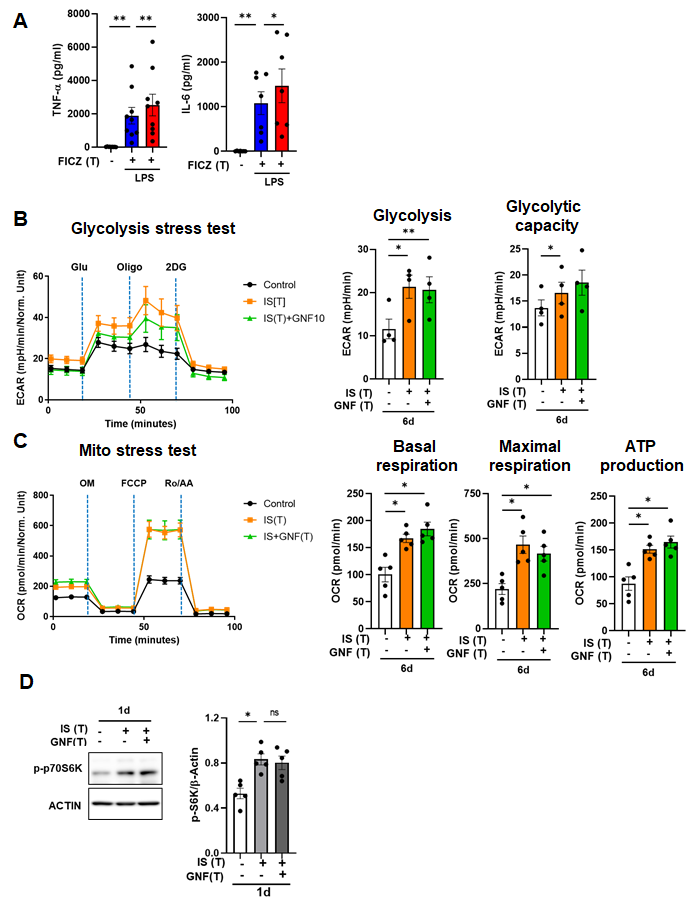

**Supplementary Figure 4. IS-mediated metabolic rewiring in IS-trained macrophages is independent of AhR. A.** Monocytes were pretreated with FICZ (100 nM), an AhR agonist, followed by training for 5 days. Cells were restimulated with LPS (10 ng/ml) for 24 hr. TNF-α and IL-6 in supernatants were quantified by ELISA. **B-C.** On day 6, IS-trained cells with or without GNF351 (10 μM) were restimulated with LPS for 24 hr. Glycolysis and mitochondrial stress test were conducted with IS-trained macrophages (n = 4 ~ 5) using Seahorse XF-analyzer. ECAR (extracellular acidification rate) levels were measured after sequential treatment with glucose, oligomycin, and 2-DG. Cellular glycolysis and glycolytic capacity were analyzed (B). OCR (Oxygen consumption rate) levels were measured after sequential treatment with oligomycin, FCCP, and Rotenone/antimycin A (Ro/AA). Basal respiration, maximal respiration, and ATP production were analyzed (C). **D**. Monocytes were pretreated with or without GNF351 followed by IS-stimulation for 24 hrs. Cell lysates were prepared and immunoblotted for phosphorylated S6K protein. Band intensity in immunoblots was quantified by densitometry. β-ACTIN was used as a normalization control. Bar graphs show the mean ± SEM. *= *p* < 0.05, **= *p* < 0.01, and ***= *p* < 0.001 by two-tailed paired *t*-test.

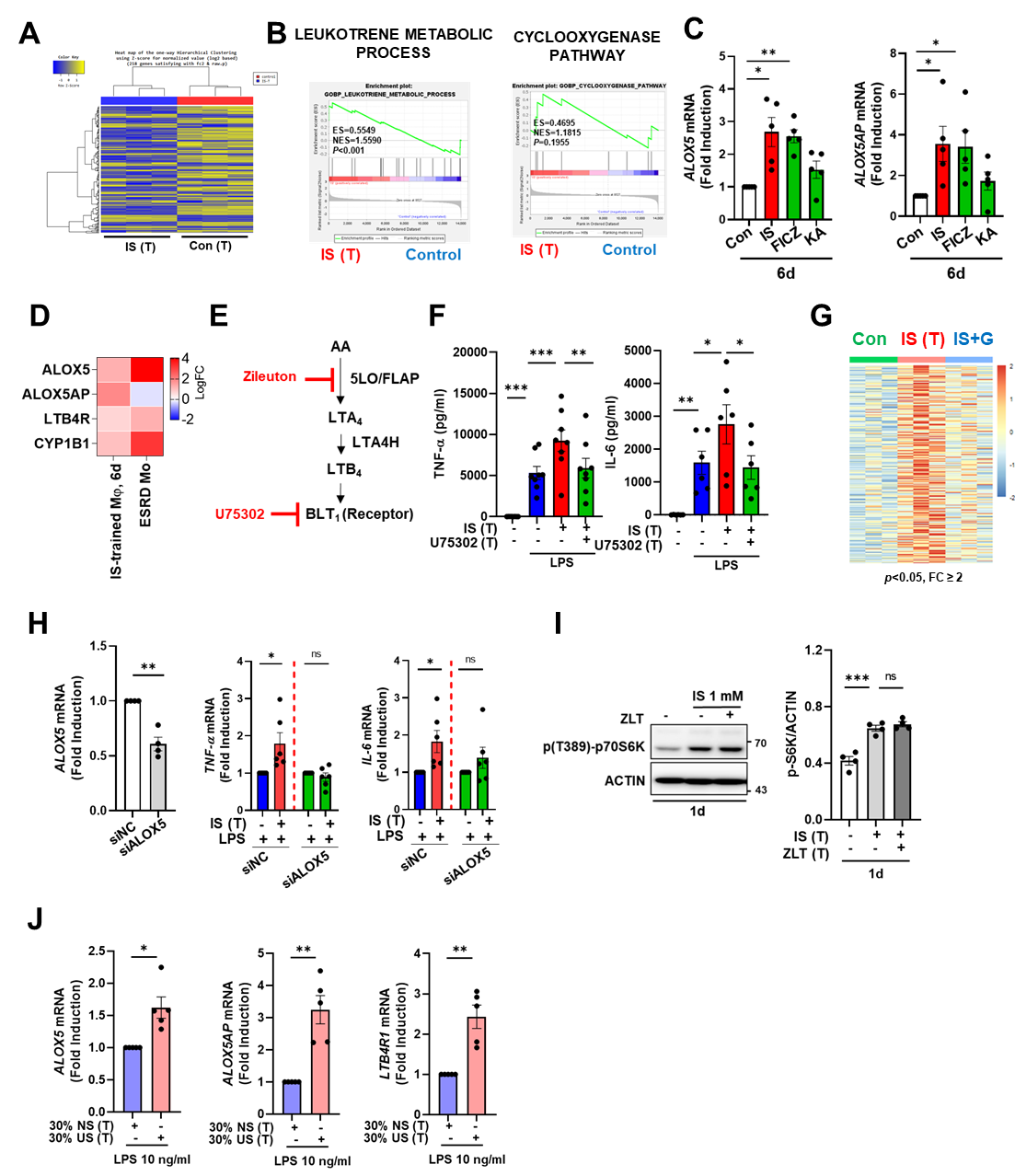
Fig. S5.

**Supplementary Figure 5.** **AhR-dependent induction of the arachidonic acid pathway contributes to IS-induced trained immunity. A.** Heatmaps of RNA-seq analysis between IS (1,000 μM)-trained and non-trained macrophages. **B.** GSEA of genes related to the leukotriene metabolic process and cyclooxygenase pathway were compared between IS-trained macrophages [IS(T)] and non-trained cells (Control). **C.** Purified monocytes were pretreated with IS (1 mM), FICZ (100 nM), or KA (0.5 mM) for 1 day, followed by 5-day resting period. mRNA expression of *ALOX5* and *ALOX5AP* was analyzed *via* RT-qPCR. **D**. Heatmaps show changes in expression of *ALOX5*, *ALOX5AP*, *LTB4R1*, and *CYP1B1* of monocytes under the indicated conditions (1^st^ lane: IS-trained macrophages, 2^nd^ lane: peripheral monocytes isolated from ESRD patients). Comparison of the fold changes of RNA-seq data in the present study and microarray data reported previously (GSE155326). **E.** Schematic diagram of the AA metabolism and target molecules of inhibitors such as zileuton and U75302. **F.** Monocytes were pretreated with U75302 (BLT1 inhibitor, 5 μM) and trained with IS for 6 days, followed by restimulation with LPS (10 ng/ml) for 24 hr. TNF-α and IL-6 in supernatants were quantified by ELISA. **G.** RNA-Seq analysis was performed on IS-trained macrophages pretreated with or without GNF351. Heatmaps of 71 upregulated DEGs including AA metabolism-related genes in IS-trained macrophages [IS(T)] compared to non-trained macrophages (Con) (Fig. 5B), illustrates their expression changes following GNF351 (10 μM) pre-treatment (IS + G). **H.** Monocytes were transfected with siRNA targeting ALOX5 (siALOX5) or negative control (siNC) for 1 day, followed by stimulation with IS for 24 hours. After a resting period of 5 days, cells were re-stimulated with LPS for 24 hours. mRNA expression levels of *TNF-α* and *IL-6* were assessed using RT-qPCR. **I.** Monocytes were pretreated with zileuton (ALOX5 inhibitor, 100 μM) and stimulated with IS for 1 day. Cell lysates were analyzed by immunoblotting. **J.** The pooled normal serum (NS) from health controls (HCs) or uremic serum (US) from patients with ESRD were used to treat monocytes isolated from HCs for 24 hr at 30% (v/v) followed by resting for 5 days. After stimulation with LPS for 24 hr, expression of *ALOX5*, *ALOX5AP*, and *LTB4R1* mRNAs were quantitated using RT-qPCR. Bar graphs show the mean ± SEM. *= *p* < 0.05 and **= *p* < 0.01, by two-tailed paired *t*-test.

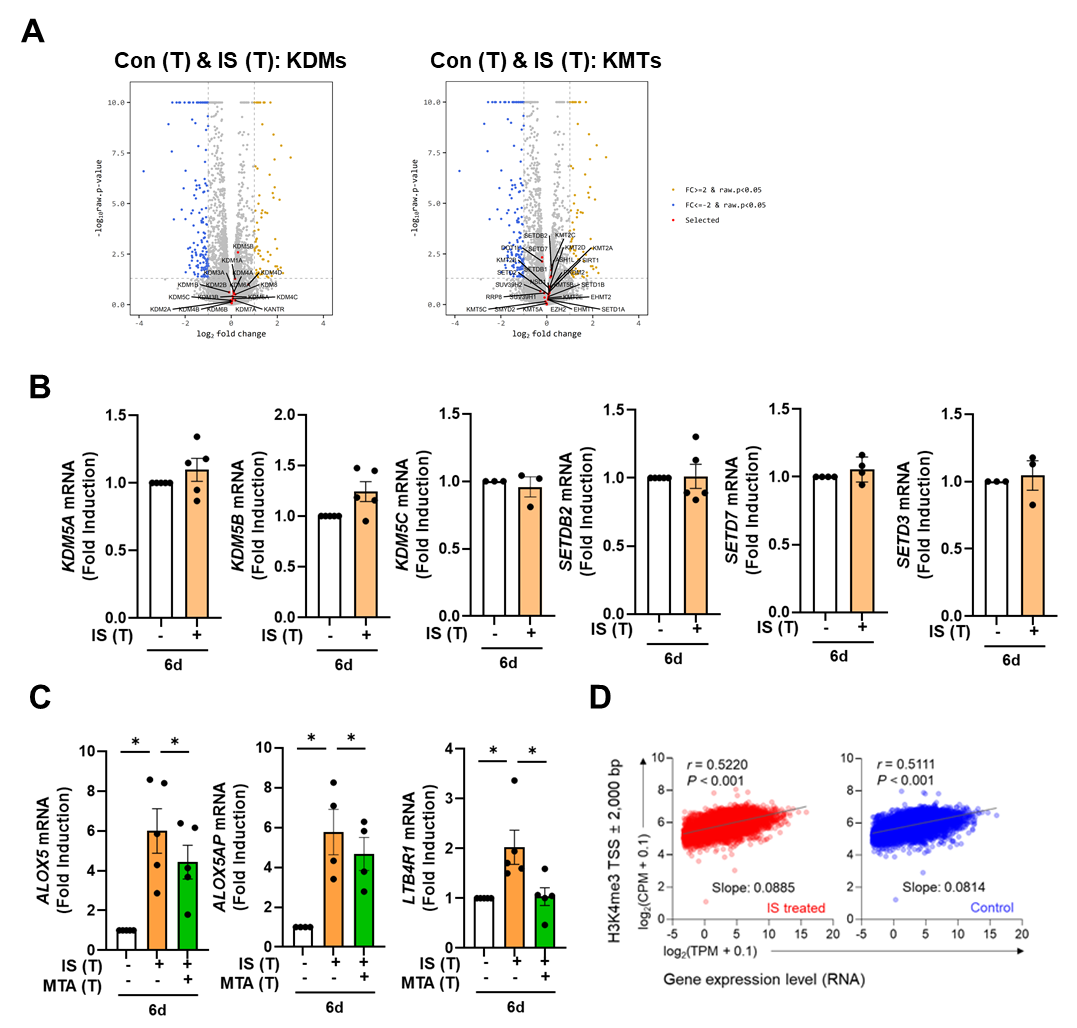
Fig. S6.

**Supplementary Figure 6. No obvious changes in expression of major histone-modifying enzymes were observed in IS-induced trained immunity. A.** RNA-Seq analysis was performed on IS (1,000 μM)-trained monocytes. Volcano plot visualized the expression of histone modifying enzymes, histone demethylases (KDMs, left plot) or histone methyltransferases (KMTs, right plot) between IS-trained and non-trained monocytes. Red dots indicate each histone modifying enzyme. **B**. On day 6 expression of *KDM5* family, *SETDB2*, *SETD7*, and *SETD3* mRNAs in IS-trained macrophages was analyzed by RT-qPCR. **C.** On day 6 after IS-training with or without MTA (200 μM), expression of *ALOX5*, *ALOX5AP*, and *LTB4R1* mRNAs were quantified using RT-qPCR. **D.** The correlation between ChIP-Seq and RNA-Seq data in IS-trained macrophages. Bar graphs show the mean ± SEM. * = p < 0.05, by two-tailed paired *t*-test.

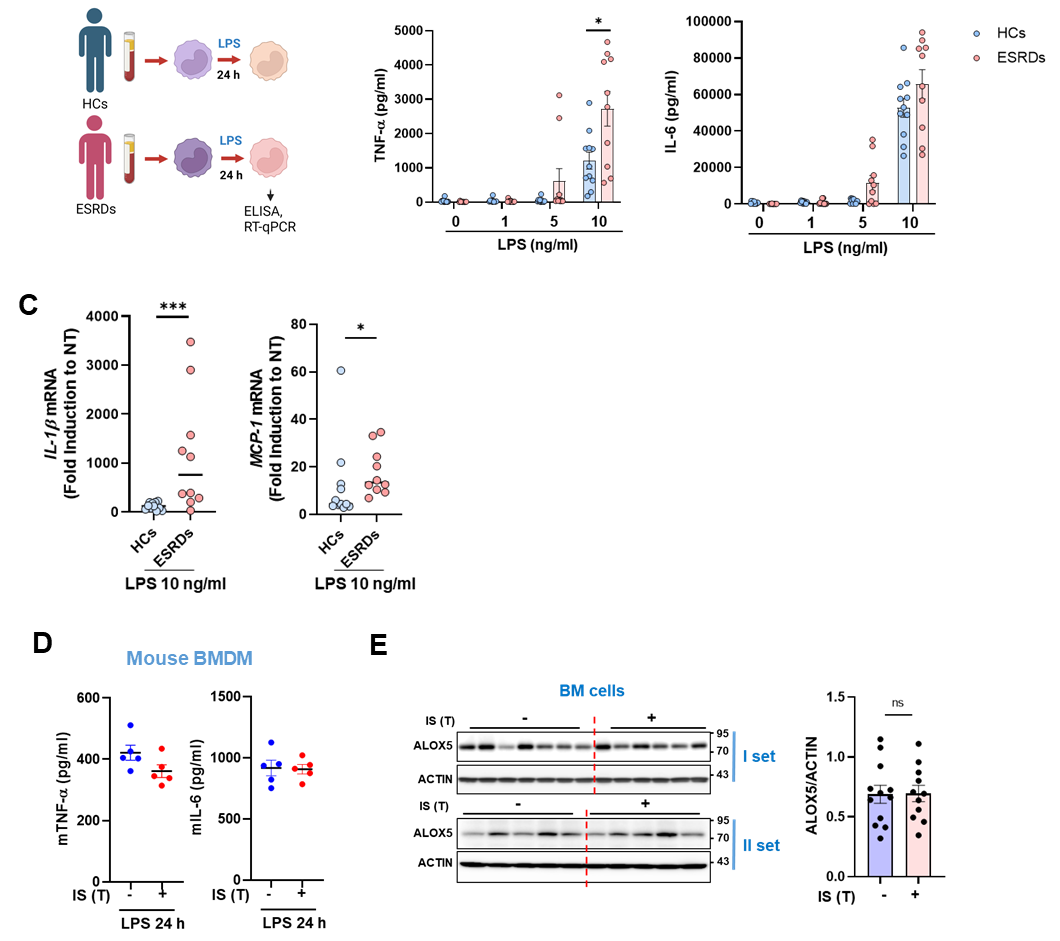
Fig. S7.

**Supplementary Figure 7. *Ex vivo* monocytes of ESRD patients exhibit features of IS-trained macrophages. A-C.** monocytes purified from ESRD patients (n = 10) and age-matched HCs (n = 11) were seeded and stimulated with LPS (10 ng/ml) for 24 hrs. TNF-α and IL-6 production were analyzed using ELISA (B) and *IL-1β* and *MCP-1* mRNA expression were determined by RT-qPCR (C). **D**. Before LPS injection, IS-trained mice were sacrificed, and bone marrow progenitor cells were mechanically separated. Isolated cells were differentiated into bone marrow-derived macrophages (BMDM) with M-CSF. On day 6, BMDM were stimulated with LPS (10 ng/ml) for 24 hours. The amount of TNF-α and IL-6 in the supernatants were quantified by ELISA. **E.** Bone marrow cells isolated from IS-trained mice were lysed. Cell lysates were prepared and immunoblotted for ALOX5 protein. Band intensity in immunoblots was quantified by densitometry. β-ACTIN was used as a normalization control. Bar graphs show the mean ± SEM (B) or the median (C). *= *p* < 0.05 and *** = *p* < 0.001 by unpaired non-parametric *t*-test.

**Table S1.** The 59 differentially upregulated enriched peaks in IS-trained cells at day 6.

| **No.** | **Fold change (IS/Ctrl)** | **Symbol** | ***P*-Value** | **chromosome** | **start** | **end** |
| --- | --- | --- | --- | --- | --- | --- |
| 1 | 2.17 | PTMA | 0.000 | chr2 | 232,572,225 | 232,572,892 |
| 2 | 2.13 | TAF9B | 0.001 | chrX | 77,394,594 | 77,395,229 |
| 3 | 1.99 | ULK1 | 0.002 | chr12 | 132,379,673 | 132,380,598 |
| 4 | 1.95 | HCN1 | 0.024 | chr5 | 46,391,617 | 46,393,029 |
| 5 | 1.92 | PRPF4B | 0.000 | chr6 | 4,018,154 | 4,019,021 |
| 6 | 1.90 | TPM2 | 0.002 | chr9 | 35,690,429 | 35,691,336 |
| 7 | 1.89 | PLCD1 | 0.008 | chr3 | 38,065,564 | 38,066,458 |
| 8 | 1.87 | ZXDA | 0.008 | chrX | 58,548,803 | 58,549,951 |
| 9 | 1.84 | CLCN5 | 0.010 | chrX | 49,683,035 | 49,684,086 |
| 10 | 1.74 | SCLY | 0.010 | chr2 | 238,968,675 | 238,969,378 |
| 11 | 1.74 | EXOSC5 | 0.015 | chr19 | 41,903,568 | 41,904,454 |
| 12 | 1.74 | ZCCHC24 | 0.012 | chr10 | 81,204,279 | 81,205,209 |
| 13 | 1.73 | NCAPG2 | 0.009 | chr7 | 158,497,836 | 158,498,379 |
| 14 | 1.66 | ZFP69B | 0.028 | chr1 | 40,889,909 | 40,890,810 |
| 15 | 1.65 | PIGP | 0.031 | chr21 | 38,442,782 | 38,443,698 |
| 16 | 1.63 | RPS12 | 0.023 | chr6 | 133,134,412 | 133,135,505 |
| 17 | 1.63 | FNBP1L | 0.027 | chr1 | 93,920,253 | 93,920,934 |
| 18 | 1.63 | KDSR | 0.036 | chr18 | 61,035,043 | 61,035,711 |
| 19 | 1.62 | MIR4436A | 0.030 | chr2 | 90,300,121 | 90,300,965 |
| 20 | 1.61 | GPSM3 | 0.033 | chr6 | 32,163,701 | 32,164,335 |
| 21 | 1.60 | FKBP11 | 0.031 | chr12 | 49,318,763 | 49,319,548 |
| 22 | 1.60 | PRKAG2 | 0.041 | chr7 | 151,605,461 | 151,606,648 |
| 23 | 1.59 | IFI16 | 0.036 | chr1 | 158,979,768 | 158,981,235 |
| 24 | 1.58 | MAOA | 0.050 | chrX | 43,514,253 | 43,514,969 |
| 25 | 1.58 | XRCC5 | 0.022 | chr2 | 216,974,068 | 216,974,992 |
| 26 | 1.58 | PQBP1 | 0.015 | chrX | 48,754,482 | 48,755,683 |
| 27 | 1.58 | TSNARE1 | 0.034 | chr8 | 143,483,367 | 143,484,264 |
| 28 | 1.57 | ENOSF1 | 0.025 | chr18 | 711,957 | 712,856 |
| 29 | 1.56 | RAD23A | 0.007 | chr19 | 13,056,634 | 13,057,623 |
| 30 | 1.56 | ACTR3 | 0.032 | chr2 | 114,646,472 | 114,647,252 |
| 31 | 1.55 | C5orf51 | 0.023 | chr5 | 41,904,386 | 41,905,346 |
| 32 | 1.55 | UCHL1-AS1 | 0.020 | chr4 | 41,258,857 | 41,260,104 |
| 33 | 1.54 | EEPD1 | 0.038 | chr7 | 36,195,035 | 36,196,312 |
| 34 | 1.54 | ZNF585B | 0.048 | chr19 | 37,700,961 | 37,701,592 |
| 35 | 1.53 | PPA2 | 0.010 | chr4 | 106,394,085 | 106,395,366 |
| 36 | 1.52 | EIF1AX | 0.016 | chrX | 20,159,079 | 20,160,075 |
| 37 | 1.51 | CD53 | 0.027 | chr1 | 111,415,818 | 111,417,201 |
| 38 | 1.51 | NUDCD3 | 0.011 | chr7 | 44,529,338 | 44,530,500 |
| 39 | 1.49 | SPATA1 | 0.012 | chr1 | 84,970,305 | 84,971,951 |
| 40 | 1.48 | HSD17B11 | 0.019 | chr4 | 88,311,038 | 88,312,383 |
| 41 | 1.47 | VPS53 | 0.042 | chr17 | 497,350 | 499,088 |
| 42 | 1.47 | FLYWCH2 | 0.045 | chr16 | 2,932,908 | 2,933,882 |
| 43 | 1.47 | RBBP9 | 0.048 | chr20 | 18,476,929 | 18,478,060 |
| 44 | 1.46 | TNFRSF21 | 0.025 | chr6 | 47,276,461 | 47,277,774 |
| 45 | 1.45 | LOC101927974 | 0.029 | chr7 | 107,384,234 | 107,385,507 |
| 46 | 1.45 | OAZ3 | 0.041 | chr1 | 151,735,094 | 151,736,365 |
| 47 | 1.44 | TMEM219 | 0.026 | chr16 | 29,973,365 | 29,974,938 |
| 48 | 1.44 | CUTA | 0.047 | chr6 | 33,384,929 | 33,386,004 |
| 49 | 1.43 | PSMA3 | 0.023 | chr14 | 58,710,630 | 58,712,355 |
| 50 | 1.43 | PLRG1 | 0.046 | chr4 | 155,470,747 | 155,472,093 |
| 51 | 1.43 | PSMA1 | 0.050 | chr11 | 14,540,951 | 14,542,589 |
| 52 | 1.40 | TMEM131 | 0.036 | chr2 | 98,611,268 | 98,612,743 |
| 53 | 1.39 | RPUSD2 | 0.030 | chr15 | 40,861,310 | 40,862,576 |
| 54 | 1.39 | NEK4 | 0.043 | chr3 | 52,803,934 | 52,805,223 |
| 55 | 1.38 | TRIP11 | 0.035 | chr14 | 92,505,410 | 92,507,100 |
| 56 | 1.37 | ACAA1 | 0.046 | chr3 | 38,177,208 | 38,178,850 |
| 57 | 1.36 | ZNF212 | 0.050 | chr7 | 148,936,596 | 148,937,656 |
| 58 | 1.35 | LRRC8D | 0.044 | chr1 | 90,286,653 | 90,288,467 |
| 59 | 1.34 | PTPMT1 | 0.048 | chr11 | 47,586,495 | 47,588,063 |

**Table S2.** Demographic characteristics in study population

|  | **ESRD (N=21)** | **HCs (N=20)** |
| --- | --- | --- |
| **Clinical variables** |  |  |
| Age (years) | 62.4±12.4 | 56.9 ±7.8 |
| Male gender (%) | 15 (71.4%) | 8 (40%) |
| CAD (%) | 6 (28.6%) |  |
| Hypertension (%) | 18 (85.7%) |  |
| DM (%) | 7 (33.3%) |  |
| SBP (mmHg) | 137.0±26.1 |  |
| DBP (mmHg) | 64.5±20.4 |  |
| Dialysis Duration (year) | 10.9±9.3 |  |
| **Laboratory variables** |  |  |
| WBC count (X 10^3^/μL) | 5.5±2.0 |  |
| Hemoglobin (g/dL) | 11.2±1.7 |  |
| Total cholesterol (mg/dL) | 152.6±37.3 |  |
| BUN (mg/dL) | 51.5±19.9 |  |
| Creatinine (mg/dL) | 8.5±3.8 |  |
| Albumin (g/dL) | 3.9±0.5 |  |
| Calcium (mg/dL) | 8.4±0.6 |  |
| Phosphorus (mg/dL) | 4.7±1.6 |  |
| hsCRP (mg/dL) | 4.9±8.6 |  |

ESRD, end-stage renal disease; CAD, coronary artery disease; DM, diabetes mellitus; SBP, systolic blood pressure; DBP, diastolic blood pressure; WBC, white blood cell; BUN, blood urea nitrogen; hsCRP, high-sensitivity C-reactive protein. Data are presented as mean ± SD or n (%)

**Table S3.** Primers for qPCR.

| **Gene name** | **Primer sequence (5’-3’)** |
| --- | --- |
| **Human *Actin*** | Forward: GGACTTCGAGCAAGAGATGG; Reverse: AGCACTGTGTTGGCGTACAG |
| **Human *TNF-α*** | Forward: TGCTTGTTCCTCAGCCTCTT; Reverse: CAGAGGGCTGATTAGAGAGAGGT |
| **Human *IL-6*** | Forward: TACCCCCAGGAGAAGATTCC; Reverse: TTTTCTGCCAGTGCCTCTTT |
| **Human *pro-IL-1β*** | Forward: CACGATGCACCTGTACGATCA; Reverse: GTTGCTCCATATCCTGTCCCT |
| **Human *IL-10*** | Forward: TGCCTTCAGCAGAGTGAAGA; Reverse: GGTCTTGGTTCTCAGCTTGG |
| **Human *MCP-1*** | Forward: AGCAGCAAGTGTCCCAAAGA; Reverse: GGTGGTCCATGGAATCCTGA |
| **Human *ALOX5*** | Forward: TCTTGGCAGTCACATCTCTTC; Reverse: GAATGGGTCCCTATGGTGTTTA |
| **Human *ALOX5AP*** | Forward: GTCGGTTACCTAGGAGAGAGAA; Reverse: GACATGAGGAACAGGAAGAGTATG |
| **Human *LTB4R1*** | Forward: GTTCATCTCTCTGCTGGCTATC; Reverse: AGCGCTTCTGCATCCTTT |
| **Human *CYP1B1*** | Forward: TGCCTGTCACTATTCCTCATGCCA; Reverse: ATCAAAGTTCTCCGGGTTAGGCCA |
| **Human *KDM5A*** | Forward: CAGCTGTGTTCCTCTTCCTAAA; Reverse: CCTTCGAGACCGCATACAAA |
| **Human *KDM5B*** | Forward: GCCCTCAGACACATCCTATTC; Reverse: AGTCCACCTCATCTCCTTCT |
| **Human *KDM5C*** | Forward: ACAGAAGGAGAAGGAGGGTAT; Reverse: CACACACAGATAGAGGTTGTAGAG |
| **Human *SETDB2*** | Forward: CCACTGAACTTGAAGGGAGAAA; Reverse: GTGGAGTGCTGAAGAATGAGAG |
| **Human *SETD3*** | Forward: TGGTTACAACCTGGAAGATGAC; Reverse: CGTTGGATCGAGTGCCATAA |
| **Human *SETD7*** | Forward: AGTGTAAACTCCCTGGCCCT; Reverse: GTTCACGGAGAAAAGAACGG |

**Table S4.** Primers for ChIP-qPCR.

| **Gene name** | **Primer sequence (5’-3’)** |
| --- | --- |
| **Human *TNF-α* promoter** | Forward: GTGCTTGTTCCTCAGCCTCT; Reverse: ATCACTCCAAAGTGCAGCAG |
| **Human *IL-6* promoter** | Forward: AGGGAGAGCCAGAACACAGA; Reverse: GAGTTTCCTCTGACTCCATCG |
| **Human *HK2* promoter** | Forward: GAGCTCAATTCTGTGTGGAGT; Reverse: ACTTCTTGAGAACTATGTACCCTT |
| **Human *PFKP* promoter** | Forward: CGAAGGCGATGGGGTGAC; Reverse: CATCGCTTCGCCACCTTTC |
